# Disease classification for whole blood DNA methylation: meta-analysis, missing values imputation, and XAI

**DOI:** 10.1101/2022.05.10.491404

**Authors:** Alena Kalyakulina, Igor Yusipov, Maria Giulia Bacalini, Claudio Franceschi, Maria Vedunova, Mikhail Ivanchenko

## Abstract

**Background:** DNA methylation has a significant effect on gene expression and can be associated with various diseases. Meta-analysis of available DNA methylation datasets requires development of a specific pipeline for joint data processing.

**Results:** We propose a comprehensive approach of combined DNA methylation datasets to classify controls and patients. The solution includes data harmonization, construction of machine learning classification models, dimensionality reduction of models, imputation of missing values, and explanation of model predictions by explainable artificial intelligence (XAI) algorithms. We show that harmonization can improve classification accuracy by up to 20% when preprocessing methods of the training and test datasets are different. The best accuracy results were obtained with tree ensembles, reaching above 95% for Parkinson’s disease. Dimensionality reduction can substantially decrease the number of features, without detriment to the classification accuracy. The best imputation methods achieve almost the same classification accuracy for data with missing values as for the original data. Explainable artificial intelligence approaches have allowed us to explain model predictions from both populational and individual perspectives.

**Conclusions:** We propose a methodologically valid and comprehensive approach to the classification of healthy individuals and patients with various diseases based on whole blood DNA methylation data using Parkinson’s disease and schizophrenia as examples. The proposed algorithm works better for the former pathology, characterized by a complex set of symptoms. It allows to solve data harmonization problems for meta-analysis of many different datasets, impute missing values, and build classification models of small dimensionality.

## 1. Introduction

### 1.1. Background

DNA methylation (DNAm) plays an important role in human development and is associated with gene expression, genomic imprinting, epigenetic modification, and other biological processes without altering the DNA sequence [1–8]. Abnormal methylation patterns can lead to numerous diseases [9]. DNA methylation consists of binding a methyl group to cytosine in the cytosine-guanine dinucleotides (CpG sites). Hypermethylation of CpG sites near the gene promoter is known to repress transcription, while hypermethylation in the gene body appears to have an opposite, also less pronounced effect. Changes in DNAm patterns are associated with aging and environmental exposures [10, 11]. Current epigenome-wide association studies (EWAS) test DNAm associations with human phenotypes, health conditions and diseases. Microarray-based technologies, such as the Illumina HumanMethylation450 (450K) and HumanMethylationEPIC (850K) arrays [12] are based on the hybridization of bisulfite-converted DNA to 50-mer probes and for each CpG site included in the design allow to estimate the fraction of methylated DNA copies. Two metrics are used to represent methylation levels: the β-value, ranging from 0 to 1, and the M-value, the log2 ratio of the intensities of methylated versus unmethylated probes [13–15]. M-values are more robust quantifiers since β-values close to 0 and 1 suffer from substantial heteroscedasticity [13].

Nowadays, machine learning has become a broadly applicable method for data modeling and analysis in a wide range of applications. The availability of large data sets and a variety of unreinforced generative methods make these approaches more accurate, simple, and relevant in bioinformatics, in particular, for transcriptomic and epigenetic data analysis [16–21]. DNA methylation data are often used for classification tasks. One of the most common examples is the classification of different types of cancer using the TCGA repository [22]. Such classifiers usually demonstrate high accuracy [21, 23–29], based on both cancer-induced changes in methylation and the differences in methylation of various tumor tissues [23, 30]. Classifying different human conditions - phenotypes or pathologies - using DNA methylation data from a single tissue is more difficult. Phenotype classification can question smoking or obesity status, although existing results suggest that such conditions may not be clearly reflected in DNA methylation [21, 31, 32]. Classification of cases and controls for certain diseases is also performed using DNA methylation data. Examples of machine learning applications using epigenetic data include classification of coronary heart disease, neurodevelopmental syndromes, schizophrenia, Alzheimer’s disease, psychiatric disorders and others [33–39].

One of the main challenges is that methylation datasets are limited in the number of samples. Increasing the amount of data requires combining many datasets collected under different conditions and then performing analysis for the merged data, which can cause a variety of problems. There are many factors that lead to significant differences in methylation data that are not directly related to the development of pathological conditions, to name the effect of the laboratory batch, different experimental conditions, normalization, and other [40].

Methylation levels can be affected by systematic variation due to biosample processing, i.e., batch-related variability (a subset of samples processed simultaneously), chip position in batches, and sample position within the chip [41, 42]. Batch effects can dramatically reduce the accuracy of measurements and produce false positive effects if the sample distribution is not uniform [43]. Most of the existing works avoid the question of the applicability of obtained models to new data. A central issue of meta-analysis is data harmonization. Ref. [44] developed an approach to systematically assess the impact of different preprocessing methods on meta-analysis. Its main advantage is the possibility of harmonization of the newly introduced datasets that does not require corrections to the previously analyzed datasets, employed for training the machine learning model.

Making use of new datasets to validate the model raises another problem, that is missing values in the data and the need to fill them in. New (test) datasets can lack information about some relatively small number of CpG sites on which the model was built. Experimental methylation data often contain missing values due to failing quality control checks, which can affect subsequent analysis. Since such missing CpG sites are necessary input parameters for the model, their values must be imputed. Examples include epigenetic clocks, which estimate biological age from small sets of pre-selected age-correlated CpG sites [5, 45–47], sensitive to small deviations in methylation levels [48]. Consequently, accurate imputation of missing data is required to improve the quality of DNA methylation analysis [49].

Dimensionality presents yet another problem. High data dimensionality is often associated with various undesirable consequences: increased computational effort, retraining, and visualization difficulties [50]. In most cases, an increase in dimensionality does not provide significant benefits, since lower dimensionality data may contain more relevant information [51]. In addition, the most common epigenetic models [5, 46, 52, 53] contain a small number of variables to simplify data processing, for better interpretation of the results and for the possibility of applying these models in real life. It is also worth noting that small DNA methylation panels are significantly less costly [54], which is an undeniable advantage for the possibility of widespread use.

Modern machine-learning-based artificial intelligence systems are powerful and promising tools [55–57]. However, while these models provide impressive predictive accuracy, their nonlinear structure makes them poorly interpretable, i.e. it is hard to explain what information in the input data leads AI to particular outputs. The need for trustworthy solutions has recently attracted much attention to methods that would “open” black box models [58–68]. This includes developing methods to help better understand what the model has learned [69, 70] as well as methods to explain individual predictions [59–61, 71, 72].

In summary, individual DNA methylation datasets contain an insufficient number of samples to apply machine learning approaches, so there is a need to combine and harmonize different datasets. Problems that arise on the way include tackling batch effects in individual datasets, missing values for certain samples, and high data dimensionality. Here, we analyze several existing fragmented solutions to these problems, develop a generalized unifying approach integrated in a pipeline, validate it and demonstrate its efficiency.

### 1.2. Study design and novelty

Our primary goal is to offer a methodologically complete pipeline for building machine learning models, classifying cases and controls for various diseases from whole blood DNA methylation data on many datasets, ranging from data harmonization to explainable artificial intelligence models. DNA methylation data are taken from different human body tissues, but the most widespread is whole blood methylation, the least invasive analysis and, therefore, of broad diagnostic prospects. We restrict our analysis to this kind of data. Our pipeline solves a problem of harmonization of methylation data from different datasets. They are collected in different laboratories, with different setups and experimental conditions. In general, the data are of different quality, and have been preprocessed differently. Harmonization is used to eliminate the unavoidable bias between the data and to minimize the associated machine learning model errors. The proposed pipeline uses harmonization with the selection of a reference dataset, in which case all other datasets are aligned with the reference one, so that when a new dataset is introduced, there is no need to renormalize the training data and hence rebuild the model. The pipeline uses the generally recognized types of machine learning models for classification on methylation data in tabular representation, in particular gradient-boosted decision trees. A hyperparametric search for the optimal combination of the parameters of these models is performed to ensure the best classification accuracy. Next, the dimensionality of the feature space is reduced to build portable models. In such models, the number of features has the same order as the most popular epigenetic models, such as the Horvath clock (353 CpG sites) [5], Hannum clock (71 CpG sites) [46], DNAm PhenoAge (513 CpG sites) [52], DNAm GrimAge (1030 unique CpGs were used to predict plasma protein levels) [53]. Such portable models allow them to be used for early diagnosis of various diseases - analysis of small CpG panels is much cheaper than full-genome analyses. Reducing the dimensionality of the data can also help discard noisy features that do not carry relevant information for classifiers. Also, the proposed approach includes the possibility of imputing missing values (CpG sites), and different approaches are used for this purpose. This is especially important when testing the model on new data, where some CpG sites critical for the model may be missed (e.g., because of technical errors in data acquisition and processing or failing quality checks). For the best models in terms of accuracy, explainable artificial intelligence (XAI) methods are applied to explore both the global influence of individual CpG sites on model predictions and to get explanations of how the methylation level values of individual CpG sites for specific subjects shape their individual predictions. Lists of the most important CpG sites in terms of machine learning models are compared with lists of CpG sites (and their corresponding genes) from existing studies associated with the considered diseases. Biological pathways of diseases based on these lists are identified and investigated.

## 2. Results

Larger training sample sizes provide better quality of machine learning models. The currently available DNA methylation data sets do not exceed several thousand samples, and that could hardly change in the near future due to complexity and cost of study. Merging different data sets, therefore, appears a practical way to circumvent size limitations. However, it poses many challenges, such as the need to harmonize datasets collected under different conditions and pre-processed in different ways, the need to fill in missing values in a way that preserves patterns in the data, the reduction of excessively high dimensionality of input variables with a relatively small number of samples. These issues have been addressed separately; below we report an integrated solution that brings together the data processing and analysis steps and the resulting methodologically complete pipeline for solving the classification problem based on merging several independent DNA methylation data.

### 2.1. Datasets and machine learning tasks

We studied whole blood DNA methylation datasets generated on subjects with Parkinson’s disease or schizophrenia. We selected 3 datasets that contain samples from subjects with Parkinson’s disease and healthy controls: GSE145361 [73], GSE111629 [74–76], GSE72774 [74, 75, 77] and 4 datasets that contain samples from subjects with schizophrenia and healthy controls: GSE152027 [78], GSE84727 [78, 79], GSE80417 [78, 79], GSE116379 (non-famine participants) [80]. Information about considered datasets is summarized in Table 1, in particular, the number of cases and controls, whether the dataset has been used as train or test, the original preprocessing type, the number of CpGs.

**Table 1.** Main characteristics of considered datasets. For each disease, the bold row represents the reference dataset for the harmonization. Number of CpGs is common for train datasets in each disease. Three largest datasets for schizophrenia, GSE84727, GSE80417 and GSE152024, have the same preprocessing.

| Disease | Dataset | Number of cases | Number of controls | Train or Test subset | Raw IDAT available? | Number of CpGs | Original preprocessing |
| --- | --- | --- | --- | --- | --- | --- | --- |
| Parkinson's disease | **GSE145361** | **959** | **930** | **Train** | **Yes** | **411761** | **Data processing: Genome Studio software** |
|  | GSE111629 | 334 | 237 | Train | Yes |  | Data processing: R software v3.4.2<br>Functional normalization: minfi R package |
|  | GSE72774 | 289 | 219 | Test | No | 411979 | Data processing: BeadStudio software v3.2 |
| Schizophrenia | **GSE84727** | **414** | **433** | **Train** | **No** | **399625** | **Importing: methylumi R package methylumiIDAT function**<br>**Preprocessing: watermelon R package pfilter and dasen functions** |
|  | GSE80417 | 353 | 322 | Train | No |  | Importing: methylumi R package methylumiIDAT function<br>Preprocessing: watermelon R package pfilter and dasen functions |
|  | GSE152027 | 290 | 203 | Test | No | 411901 | Importing: methylumi R package methylumiIDAT function<br>Preprocessing: watermelon R package pfilter and dasen functions |
|  | GSE116379 | 51 | 54 | Test | No | 407781 | Removed: X and Y chromosome, non-specific binding probes, failed probes based on a detection p-value larger than 0.001 and bead count lower than 5, probes with SNPs of Minor Allele Frequency larger than 5 percent within 10 base pairs of the primer<br>Functional normalization: minfi R package |

For each disease, we built machine learning models to classify cases vs. controls. Some of these datasets are used as train data for building the model, and the rest is used to test the model. For each disease we selected a reference dataset, against which harmonization was performed. As it can be seen from Table 1, the original preprocessing is the same for the majority of the considered datasets with schizophrenia patients, but it varies considerably among the different datasets for Parkinson’s disease. To reduce the influence of the laboratory-specific data collection and processing conditions on classification results, harmonization is necessary.

### 2.2. Meta-analysis and harmonization

Combining different DNA methylation datasets can improve the statistical power to test hypotheses and identify epigenetic signatures by meta-analysis. However, such meta-analysis also poses serious problems related to data harmonization, which is often not considered [81]. This is especially true for DNA methylation, where data is often only available in the preprocessed rather than raw form, and where diverse preprocessing pipelines are used [44]. Developed in [44] approach regRCPqn (regional regression on correlated probes with quantile normalization) allows for meta-analysis even if the raw data are not available. Importantly, as emerging datasets are aligned, the already treated datasets do not require renormalization. Therefore, we apply this approach to harmonization with reference. The largest dataset for each disease is taken as the reference, and other datasets are harmonized relative to it. The schematic representation of the harmonization process is shown in Figure 1.

**Figure 1.**
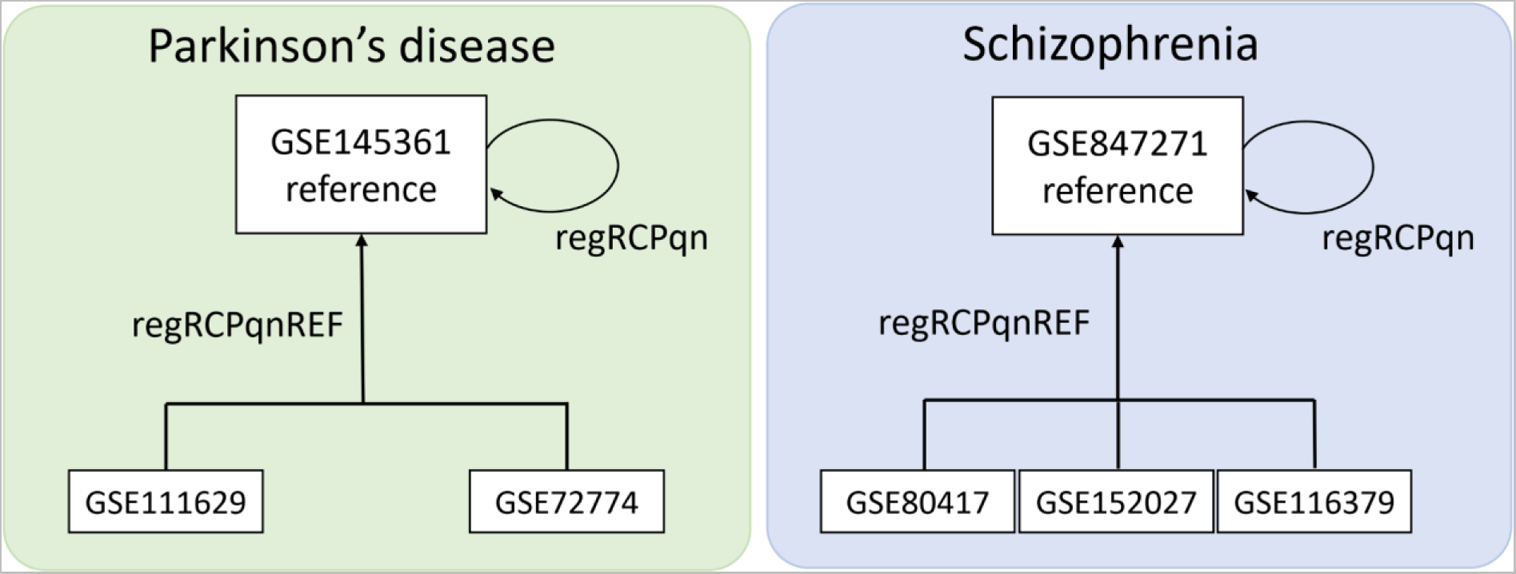
Schematic representation of harmonization procedure for Parkinson’s disease (left) and Schizophrenia (right) datasets.

For machine learning models, we used only those CpG sites that have the same distribution of methylation levels in different datasets in the control group (methylation levels in the case group typically have greater variability because of disease heterogeneity). We used the Mann-Whitney U-test [82] to compare DNA methylation values of healthy participants from the considered train datasets before and after harmonization. After harmonization, the number of CpG sites with the adjusted p-value >0.05 (not significantly different between healthy subjects from the considered train datasets) increased from 43019 to 50911 for Parkinson’s disease and from 35145 to 110137 for schizophrenia. Figure 2 illustrates the change in the distributions of methylation level values before and after harmonization. In particular, CpG sites whose methylation level distributions differed significantly before harmonization (FDR-corrected p-values<0.05) manifest similar distributions after harmonization (FDR-corrected p-values>0.05).

**Figure 2.**
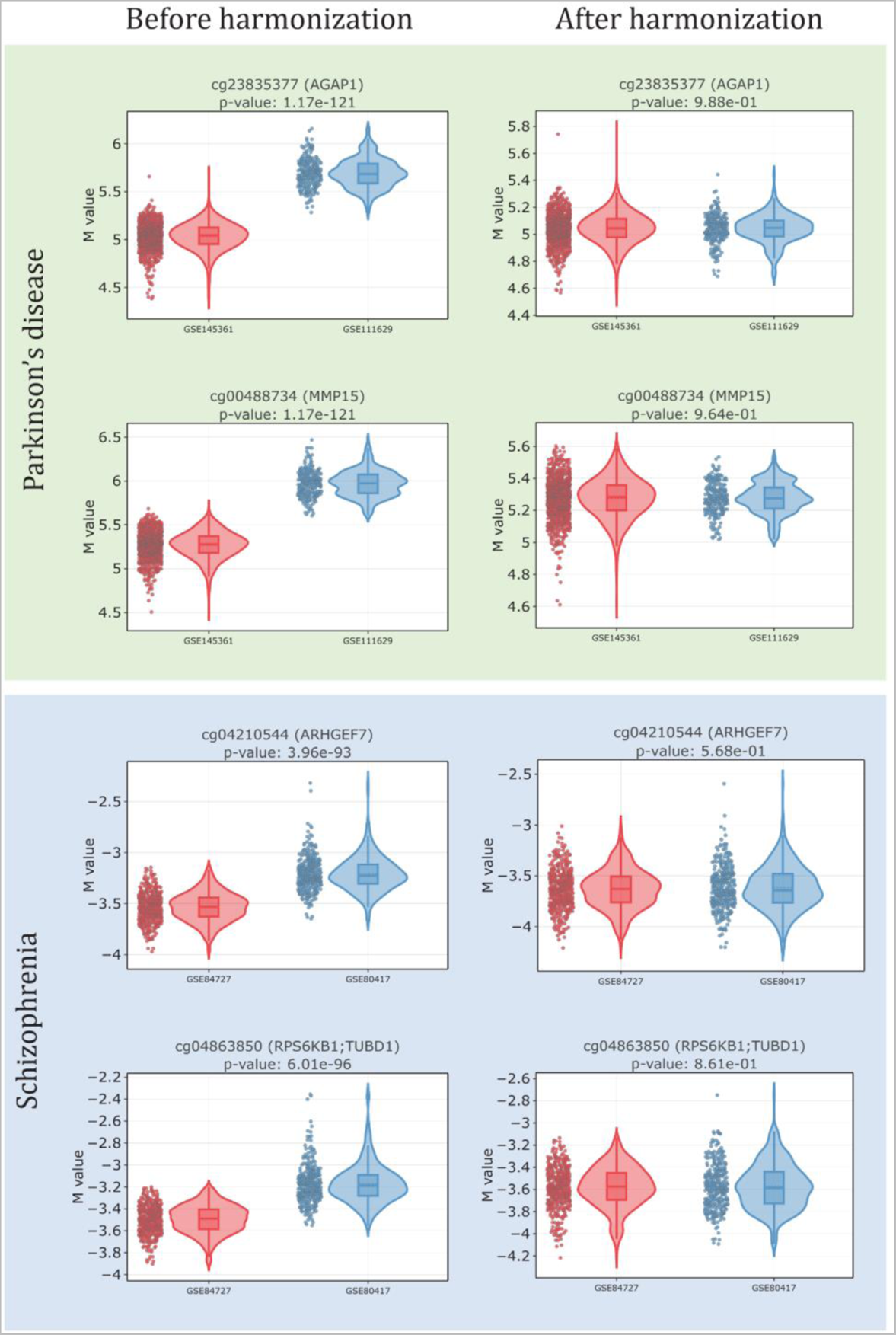
Examples of M-values methylation levels distribution for control groups before (left column) and after (right column) harmonization for Parkinson’s disease (green background) and Schizophrenia (blue background).

### 2.3. Classification models

The most common type of data representation for machine learning is tabular, and DNA methylation data fulfills it. Typically, the rows refer to participants, the columns refer to CpG sites, and the cells of the table contain the methylation levels of each CpG site for each participant. There are many machine learning models designed to work with tabular data: XGBoost [83], CatBoost [84], LightGBM [85], TabNet [86], NODE [87]. Main characteristics of the models are summarized in Table 2.

**Table 2.** Main characteristics of the considered classification models.

| Model | Type | Feature importance API | Missing values imputation |
| --- | --- | --- | --- |
| XGBoost | Gradient-boosted decision tree ensemble | Yes | Learned during training |
| CatBoost | Gradient-boosted decision tree ensemble | Yes | Min or Max for numerical features |
| LightGBM | Gradient-boosted decision tree ensemble | Yes | No information |
| TabNet | Deep neural network | Yes | Mean for numerical features, Vector of zeros in one-hot encoding for categorical features |
| NODE | Gradient-boosted decision tree ensemble | No | No information |

For each disease, all considered datasets were divided into training and test ones (as stated in Table 1). We trained all models on two training datasets and then tested on the remaining datasets. Accuracy with weighted averaging was the main quality metric, as it can handle situations with possible imbalance of the classes (the number of participants in different classes varies significantly). As discussed in the above, the approach fulfills the requirement that the model does not have to be trained again as the new data set is considered. Moreover, the models must be trained to classify biological differences in methylation data rather than traces of different experimental conditions in different laboratories. Accordingly, we do not mix train and test datasets and do not perform cross-validation.

Newly introduced datasets may lack some CpG sites that are present in already trained models; in this case various imputation methods are applied (cf. Sections 2.5, 4.5 for more details). Models for Parkinson’s disease for non-harmonized data are trained on 43019 CpG sites, for harmonized data on 50911 CpG sites. Among these, the Parkinson’s disease test dataset GSE72774 lacks 38 CpG sites in the non-harmonized data and 34 CpG sites in the harmonized data. Models for schizophrenia for non-harmonized data train on 35145 CpG sites, for harmonized data train on 110137 CpG sites. The first test dataset for schizophrenia GSE152027 lacks 9 CpG sites in the non-harmonized data and 36 CpG sites in the harmonized data. The second test dataset for schizophrenia GSE116379 lacks 268 CpG sites in the non-harmonized data and 609 CpG sites in the harmonized data. These values are also shown in Figure 3.

**Figure 3.**
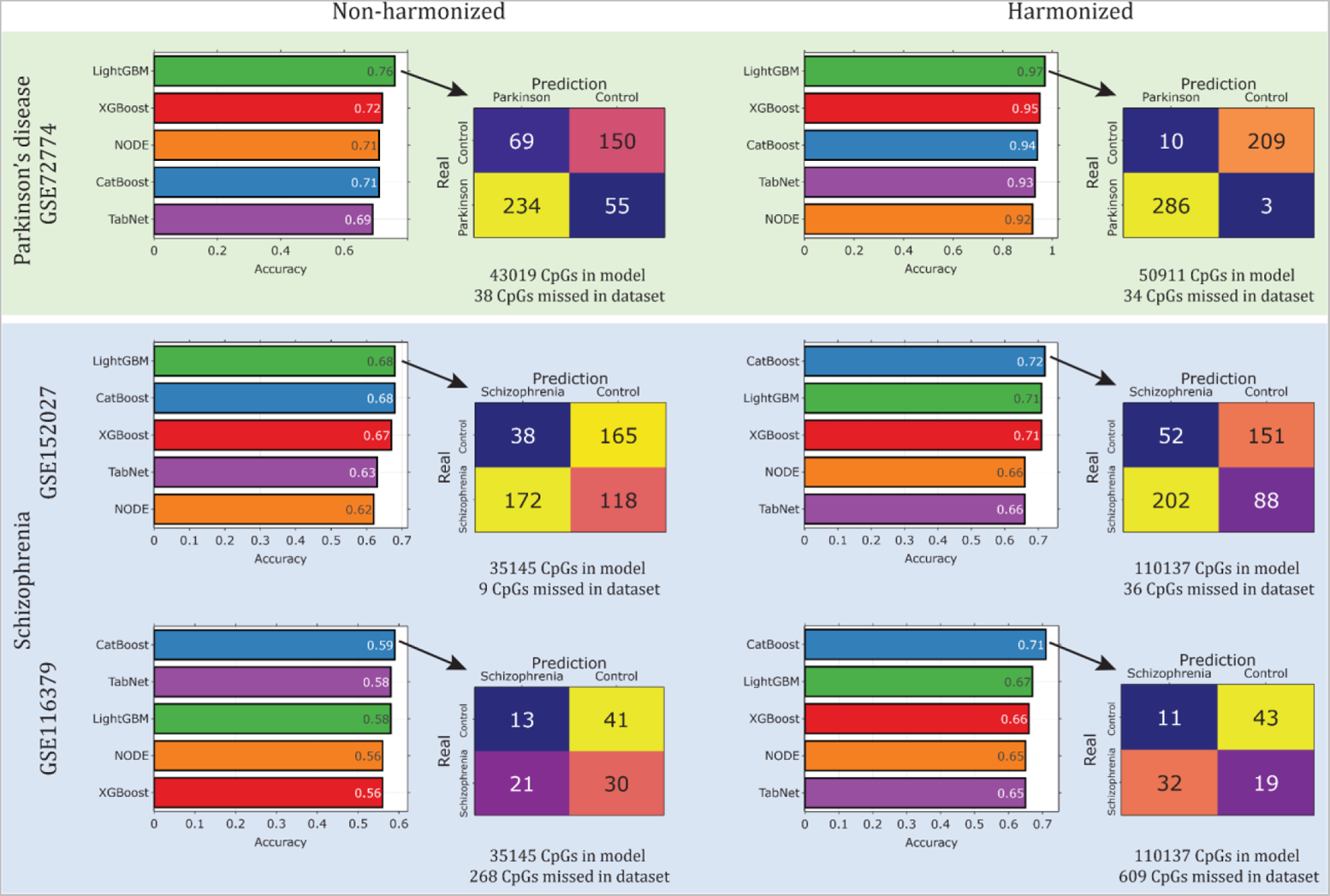
Binary classification results of baseline models for non-harmonized (left) and harmonized (right) data. For Parkinson’s disease (green background) and schizophrenia (blue background), results comparing the accuracy of different models for non-harmonized and harmonized methylation data are shown. For each test dataset, a barplot with a ranking of the weighted accuracy of the different models is shown. A confusion matrix is presented for the best model for each experiment, indicating the number of correct and incorrect predictions for each class. For each experiment, the total number of CpG sites on which the models were trained, as well as the number of missed CpG sites, are also shown.

Figure 3 shows the results of cases vs. controls classification by baseline models based on non-harmonized and harmonized whole blood methylation data for Parkinson’s disease and schizophrenia on test datasets. For each combination of harmonization type, disease, and test dataset, the barplot with the best weighted accuracy values for all constructed models is given. Confusion matrix showing the number of correct and incorrect predictions is also given for the best model (in terms of weighted accuracy). For each disease, the number of CpG sites on which the models were trained is shown, as well as the number of missing values in each experiment. All imputation methods described in Section 2.5 do not significantly change the quality of the resulting models, because all missed CpGs in test datasets have a very low value of feature importance in models with corresponding API.

The results confirm that harmonization must be applied and is most efficient for the datasets with different preprocessing methods. In particular, for Parkinson’s disease, all datasets have different original preprocessing, and the best model trained on such data shows a result of 76%. When these data are harmonized, accuracy improves dramatically to 97%. For both non-harmonized and harmonized data, for Parkinson’s disease, the best model in terms of weighted accuracy is LightGBM. For schizophrenia, only one of 4 datasets has a different preprocessing (GSE116379, Table 1). Then, harmonization does not significantly affect the quality of the built models if the datasets have the same preprocessing (68% without harmonization, 72% with harmonization in the best models). The best model is LightGBM for non-harmonized data and CatBoost for harmonized data. However, applying models trained on non-harmonized data to data with a different preprocessing gives a poor result for binary classification - 59%. Harmonization of data improves the performance of the trained models, making them close to the best results obtained for schizophrenia in terms of quality - 71%. It is also worth noting that the overall classification quality for these two diseases on methylation data is very different, possibly due to the different etiology and molecular mechanisms involved in the two diseases.

Best accuracy models allow us to extract importance values for all features. The ranking of the most important features for these models for Parkinson’s disease and schizophrenia is shown in Figure 4. It is worth noting that for schizophrenia, there is one outstanding CpG with the highest importance for classification, while the others have much lower values. For Parkinson’s disease, the situation is more uniform. These rankings can be used for the dimensionality reduction of the built models.

**Figure 4.**
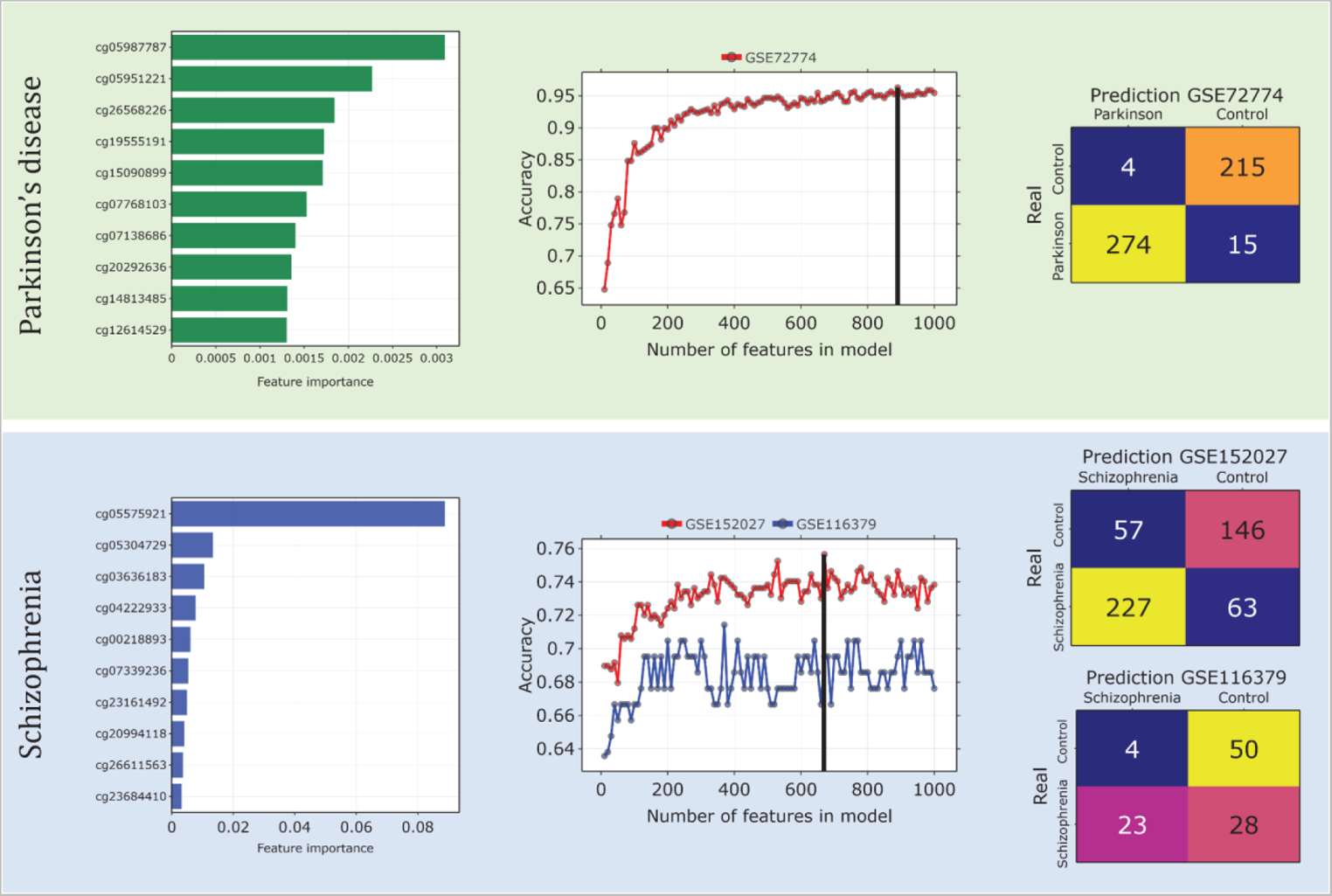
Dimensionality reduction for the best classification models for Parkinson’s disease (green) and Schizophrenia (blue). Left barplots represent top-10 features for the best classification models with the normalized importance values. Center plots show the dependence of the weighted accuracy on the number of features in the model. Black lines mark the optimal small models, consisting of 890 CpG sites for Parkinson’s disease and 670 CpG sites for Schizophrenia (GSE152027). Confusion matrices to the right show predictions for different test datasets.

### 2.4. Dimensionality reduction

As a result of applying different baseline models to methylation data to classify cases vs controls, the ones with an API for feature extraction showed the best accuracy. Based on the obtained ranking of the features (Figure 4), we performed dimensionality reduction of the constructed models. Most common epigenetic models comprise few CpG sites, no more than several hundred (for example, those used to predict epigenetic age like Horvath’ clock and Hannum clock). First, models based on few features show significantly better performance while maintaining similar classification accuracy. Second, such models are less memory-consuming.

Along these lines, we reduced the dimensionality of the model, leaving only the most important features for classification. Figure 4 shows the dependence of weighted classification accuracy on the number of features in the model for Parkinson’s disease and for schizophrenia. It first increases, until reaching a certain optimal number of features, and then changes weakly. For Parkinson’s disease, the best weighted accuracy of 96% is observed for 890 CpG sites, with an accuracy value changed by only 1% compared to the full data (50911 CpG sites). For schizophrenia, the best weighted accuracy of 75% is observed for 670 CpG sites for the test dataset GSE152027 and 70% for the test dataset GSE116379. The optimal model for schizophrenia was chosen as the one for the GSE152027. The accuracy values changed by no more than 3% compared to the full data (110137 CpG sites). The list of CpG sites that make up these small models, as well as basic information about them (gene, chromosome, relation to the CpG island) is presented in Supplementary Table S1. The resulting CpG lists were compared with previously published lists of biomarkers associated with Parkinson’s disease [74, 88, 89] and schizophrenia [78, 90]. Interestingly, for Parkinson’s disease, there is practically no overlap with the previous results, except for one CpG site from [88]. This CpG belongs to the gene DYNC1H1, which is associated with neurological and neurodegenerative diseases [91, 92]. For schizophrenia, only 15 CpG sites are common with [78]. Some of the connected genes like PRKCZ, SHANK2, ZNF608, PRDM16 were also identified as schizophrenia risk factors [93–96]. Genes, corresponding to CpG sites, from the optimal small model for Parkinson’s disease were enriched in several gene ontologies related to neuronal and metabolic processes, whereas genes from schizophrenia models were enriched in gene ontologies related to cell development processes (Supplementary Table S2).

### 2.5. Imputation of missing values

For trained machine learning models (both large and small), it is important that there are no missing values in the upcoming test data. Since it is impossible to guarantee their absence, various imputation methods are used to fill them in. Not all models support data imputation, and since some of them support only a limited set of imputation methods (Table 2), we use several of them: mean, median, mode, random, chained equations, expectation maximization, KNN with different numbers of neighbors (from 1 to 3). To study the effect of these imputation methods on the classification accuracy, we consider the following simulation experiment. For each disease, we “remove” 100 CpG sites with the highest importance values and impute them. The number of CpG sites was chosen to induce a significant drop in accuracy and to sharpen the differences in efficiency between the imputation methods. The missing CpG sites shown in Figure 3 do not take part in the construction of small optimal models, so the actually existing CpG sites are removed from consideration. Figure 5 shows the barplots for the test datasets under consideration. For Parkinson’s disease, KNN with one neighbor kept the classification accuracy at the baseline level of the data without missing values. Imputation with mode also showed good results, losing only 3%. The other methods achieved an accuracy of no more than 90%. For schizophrenia, none of the approaches achieved the baseline accuracy for data without missing values. This may be explained by the critical importance of specific features for classification. KNN with one neighbor for both datasets shows one of the best imputation results, for GSE152027 chained equation and expectation maximization perform better than KNN by 3% and 2%, respectively. Median and random values methods show unsatisfactory results in all experiments.

**Figure 5.**
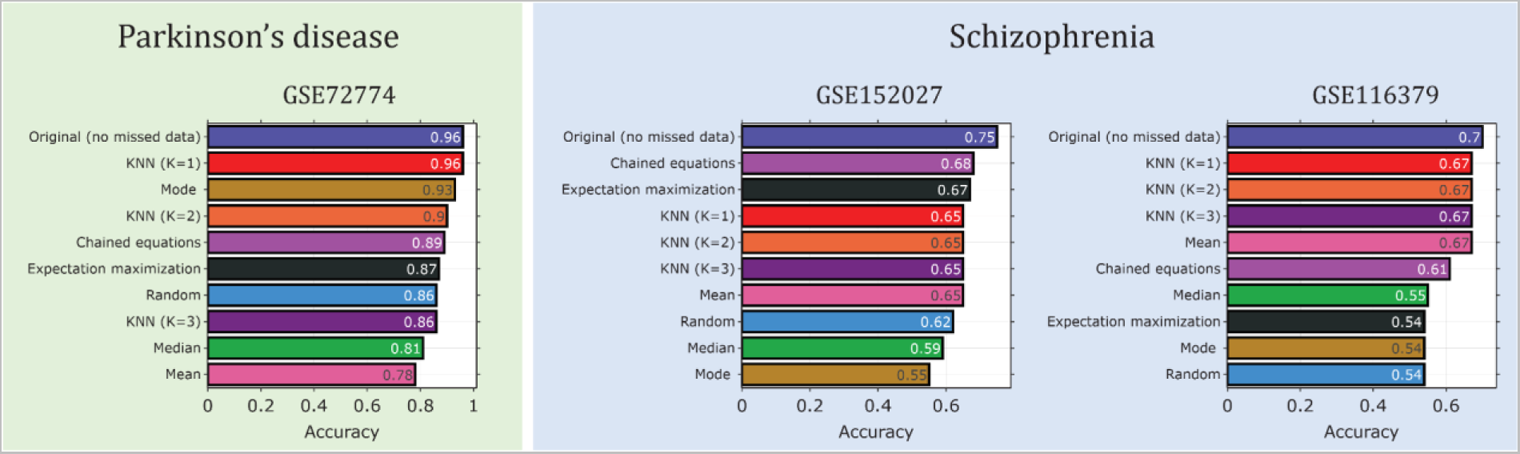
Comparison of different missing value imputation methods and their effect on weighted classification accuracy for Parkinson’s disease (green background) and schizophrenia (blue background). In all cases, 100 CpG sites with the highest importance values were dropped.

### 2.6. Explainable artificial intelligence

Even the most accurate machine learning models make mistakes on upcoming data. It presents a major challenge for those models that work as “black boxes” with unknown principles behind made decisions. SHapley Additive exPlanations (SHAP) help to understand why the model makes its predictions from the global and local points of view [97].

The global explainability of the constructed models on the training data for Parkinson’s disease and schizophrenia is illustrated in Figure 6. Beeswarm plots show the relationship between SHAP values and methylation levels for the most important CpG sites. For each CpG site, the distributions of methylation levels for all participants are shown. In particular, for Parkinson’s disease, most of the participants in the CpG site cg05987787 have low methylation levels, which positively affects the probability of predicting the disease. The right scatter plot shows in detail the distribution of methylation levels in different participants and the SHAP value. We can see that M-values below 4 have a positive effect on the probability of predicting disease, while M-values above 4 have a negative effect. The black line divides the areas of positive and negative influence of SHAP values on the prediction of disease probability. The opposite situation is observed for the CpG site cg05951221. M-values below −1 have a negative effect on the probability of predicting disease, while M-values above −1 have a positive effect. Similar plots are shown for schizophrenia. The beeswarm plot shows that there is one most important CpG site, cg05575921, that contributes the most to the probability of predicting disease, as previously shown, and the other CpG sites have a much smaller effect.

**Figure 6.**
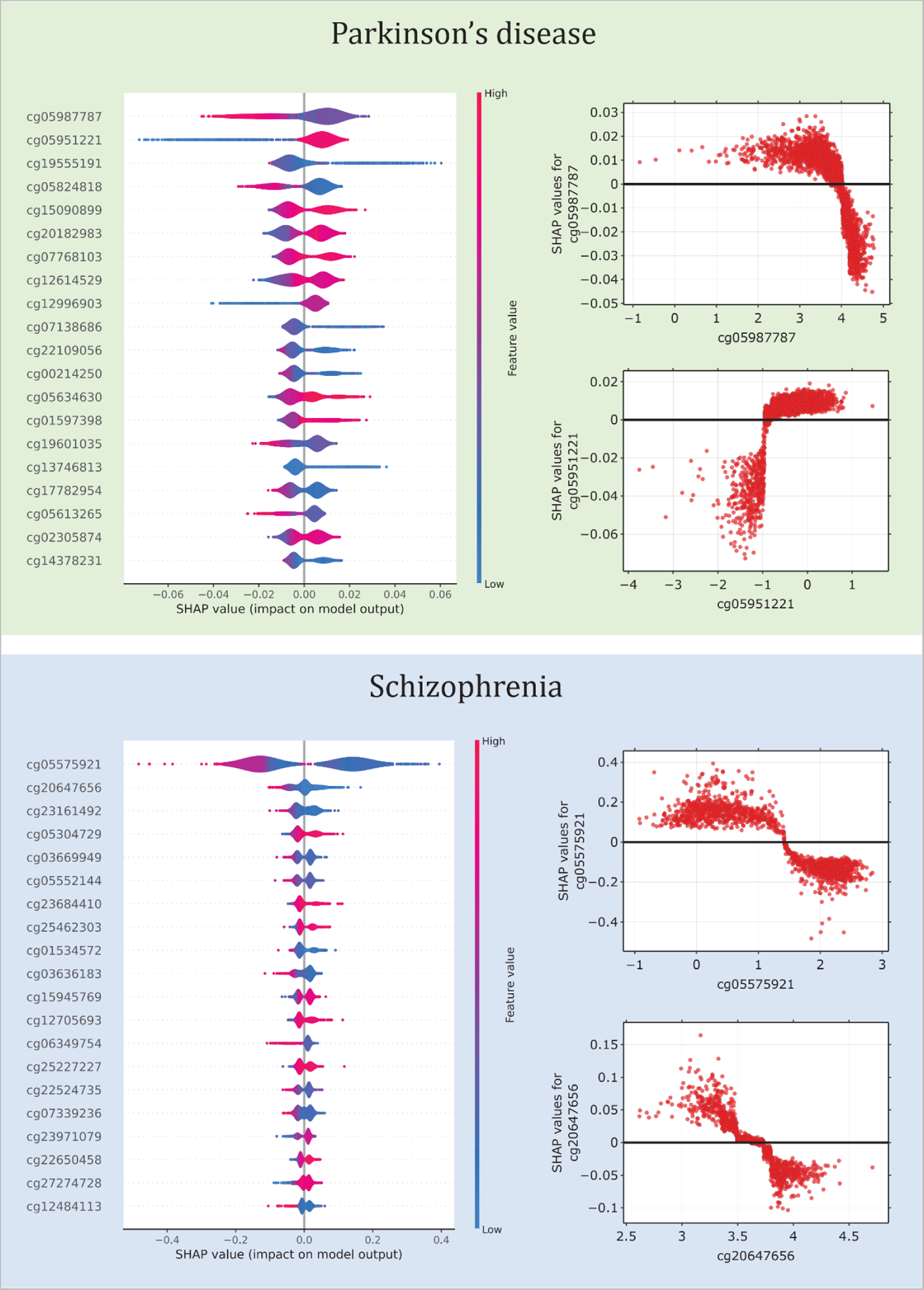
Global explainability based on SHAP values for Parkinson’s disease (green background) and schizophrenia (blue background) on training data. (Left) Beeswarm plots show the dependence of SHAP values for each CpG site on their methylation levels. Each dot represents one participant. (Right) Examples of the dependence of SHAP values on M-values methylation levels for individual CpG sites. The black line separates the areas of negative and positive influence of SHAP values on the probability of predicting the corresponding disease.

The local explainability of predictions on the test data is shown in Figure 7. The top row presents heatmaps with participants on the x-axis, CpG sites on the y-axis, and SHAP values encoded on a color scale. The participants are ordered based on the probability to predict the disease. Model output is shown above the heatmap matrix. The black line represents the probability of predicting the disease for each participant. It follows that for Parkinson’s disease, where the model works with high accuracy, this line is quite smooth and similar to the softmax function. Whereas for schizophrenia, for which the models have much lower accuracy, these probability plots are more fragmented. As shown earlier, for schizophrenia, one CpG site is the most important, so it has the highest absolute SHAP values and appears the brightest in the heatmaps. The center and bottom lines represent waterfall plots for participants with the disease and controls, respectively. They allow for explaining the model output for each participant separately. The bottom part of the waterfall plot shows the base probability of the model to predict disease, and then each line shows how a positive (red) or negative (blue) contribution from each CpG site moves the probability to the model output for that prediction. The output of the model is the probability of predicting disease. If the probability is greater than 50%, the model identifies the participant as a case, otherwise it identifies the participant as a control. The baseline probability is the average probability of the model predicting on the test data. Because baseline probability is a characteristic of the model, it depends on the quality of the model. If the model has reasonably good accuracy, then the base probability is close to the proportion of participants in a particular class. In the examples from middle line of the Figure 7 for all participants with diseases, the probability of predicting disease in the examples was above 97% (models identify them as cases almost for sure); for control participants (bottom line of Figure 7), the probability of predicting disease was below 5% (models identify them as controls almost for sure).

**Figure 7.**
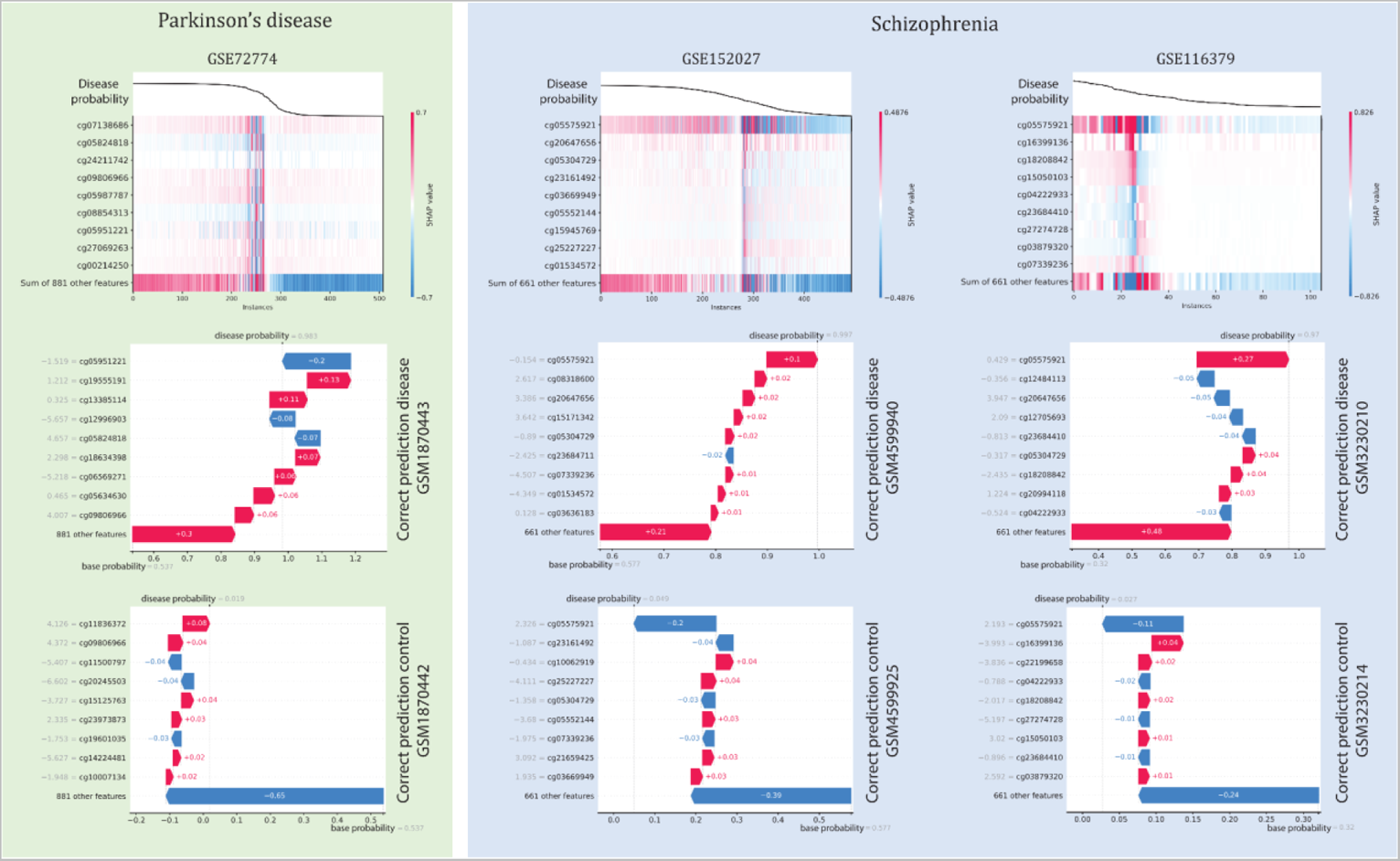
Local explainability based on SHAP values for Parkinson’s disease (green background) and schizophrenia (blue background) on test data. (top line) Black lines show the probability of predicting disease for different participants. Heatmaps in color show the contribution of SHAP values to the probability of predicting disease for different participants and CpG sites. (center line) Waterfall plots for participants with disease, showing the contribution of individual CpG sites to changes in the probability of predicting disease. In all cases, the models’ confidence is over 97%. (bottom line) Waterfall plots for control participants showing the contribution of individual CpG sites to the change in probability of predicting disease. In all model cases, the probability of predicting disease is below 5%. All waterfall plots have a member code from the GEO repository.

## 3. Discussion

### 3.1. Conclusion

We developed a multifunctional pipeline for applying machine learning models to classify cases and controls for different diseases based on DNA methylation data. Specifically, we considered Parkinson’s disease and schizophrenia as examples of complex diseases requiring early diagnosis. In addition, for these diseases there are large publicly available whole blood DNA methylation datasets. The first step of the pipeline is to harmonize the data according to the chosen reference dataset. We have shown that harmonization works well also when the available datasets were preprocessed using different pipelines and tools, as it often occurs. Harmonization can increase the classification accuracy by up to 20%. This is fully consistent with the original paper, which proposed the harmonization method regRCPqn [44]. When the preprocessing of training and test data is the same, harmonization has almost no effect on the final classification accuracy. It is impossible to guarantee that all new datasets on which the model will be tested will have the same preprocessing. Even with tools such as limma [98], ComBat [99], it may not be possible to remove technical signal when batches are mixed with variables of interest. Applying ComBat to high-throughput data with an uneven study design may actually result in false signals [100]. To solve the classification problem, different models were tested, both gradient-boosted decision trees and deep neural networks. For all models, a hyperparametric search was performed to select the optimal set of parameters. The best results of weighted accuracy were obtained with tree ensembles, which is consistent with other works [101]. For Parkinson’s disease, the accuracy of classifying patients and healthy controls was higher than 95%. The accuracy for schizophrenia was much lower, only >70%. Since the best models allow us to obtain the importance values of each feature for classification, we constructed a CpG site ranking for each of the diseases. Based on this ranking, we reduced the dimensionality of the classification models, since many of the most common epigenetic models (such as the epigenetic clocks) contain relatively few CpG sites (up to 1000). To find optimal small models, we performed a series of experiments for an increasing number of features from 10 to 1000. Based on the dependence of the weighted classification accuracy on the number of features, we determined the optimal models, with the accuracy for both diseases changing not much, while the number of features decreased significantly. Optimal small models contain 890 CpG sites for Parkinson’s disease and 670 CpG sites for Schizophrenia. Such small models are much less memory-consuming and have better computational performance. For machine learning models (both small and large), missing values are critical. A model trained on a particular set of features must necessarily be tested on the same set. If, for certain reasons (data collection or processing errors, low signal intensities, or other problems), some CpG sites are missing from the new data for testing, imputation methods are applied. The best imputation methods recover almost the same classification accuracy as for the original data. Since machine learning models work as a “black box”, special methods must be used to explain exactly how the model makes predictions. Otherwise, the model cannot be trusted and it is impossible to identify the nature of errors. If the predictive capabilities of the model can be explained, it can help to discover complex relationships between biomarkers. The calculation of SHAP values allowed us to obtain both global and local explainability. Globally, the prediction of healthy control or patient with Parkinson’s disease is affected by several CpG sites, there are examples of both types of influence, positive and negative. Global explainability for healthy controls and patients with schizophrenia confirmed the strongest influence of one CpG site, cg05575921, while the others had little effect on model prediction. This CpG site has previously been reported to be associated with schizophrenia and post-traumatic stress disorder (PTSD) [79, 102]. Examples of local explainability - specific predictions for certain participants - are also given.

Thus, we propose a methodologically valid and complete approach for classifying healthy people and patients with various diseases, which allows to harmonize DNA methylation data from different sources, impute missing values, reduce dimensionality of the models, and apply explainable artificial intelligence approaches. The proposed algorithm works better for Parkinson’s disease than for schizophrenia, which is characterized by a variety of different symptoms. Further work may include expanding the pool of considered diseases, enriching the library of methods at different stages of the pipeline. Another future challenge is advancing from explainability to interpretability. The former, currently implemented, uncovers the internal “mechanics” of a system. The yet missing interpretability would predict the outcome of changing the input or algorithmic parameters.

### 3.2. Limitations

The proposed approach has several limitations relevant at different steps. First, it should be noted that the accuracy gain from harmonization will be limited if the preprocessing of training and test data is the same. Second, the constructed classification models may not be globally optimal in terms of quality metrics, because we are considering a limited number of parameters to vary within a hyperparametric search, each varying within a limited range of values near defaults. Third, besides choosing the top best features to reduce dimensionality, it might be better to consider different combinations of them. However, in this case, the number of considered models would increase dramatically. Fourth, using schizophrenia as an example, it was shown that if there are some most important CpG sites significantly overcoming the importance of the other, and if they are missed, imputation can not help to improve the result.

## 4. Methods

### 4.1. Datasets

We reviewed publicly available whole blood DNA methylation datasets in the GEO repository [103], which include the largest ones from patients with Parkinson’s disease and schizophrenia, with at least 50 participants in each group. The following datasets comprise whole blood samples from subjects with Parkinson’s disease and healthy controls: GSE145361 [73], GSE111629 [74–76], GSE72774 [74, 75, 77]. The following datasets comprise whole blood samples from subjects with schizophrenia and healthy controls: GSE152027 [78], GSE84727 [78, 79], GSE80417 [78, 79], GSE116379 (only non-famine participants) [80]. We remove from the analysis non-CpG probes [104], SNP-related probes [105], multi-hit probes [106], probes on chromosomes X and Y. We consider only common CpGs in train datasets, the number of CpGs in test datasets can be different.The remaining amount of CpGs after all filtration procedures is shown in Table 1.

### 4.2. Data harmonization

Meta-analysis can be done in different ways: some approaches first analyze different datasets separately and then combine the results into a final estimate; others first combine data from all sets and then analyze the combined data using a single model. The first class of approaches includes aggregated data and two-step meta-analysis of individual participant data (IPD) [107]. The major advantage of these approaches is their relatively low implementation complexity, while the major disadvantage is the need for raw data. This approach often uses not all available data, but only a subset (usually differentially methylated positions, DMPs, or differentially methylated regions, DMRs). The second class of approaches is called single-step IPD meta-analysis. Although single-step IPD approaches are expected to behave similarly to two-step IPD [107], they provide additional flexibility (e.g., no need to start with raw data) and enable comparison between different models [107]. An important assumption of single-step IPD meta-analysis is the comparability of variables measured in different datasets [108], and therefore data harmonization is crucial to ensure that methylation samples of the same type (same tissue, health status, age, gender, etc.) from different datasets can be compared.

We use the approach to data harmonization proposed in [44], a one-step IPD approach for systematically assessing the impact of different preprocessing methods on the meta-analysis. It has been shown that data preprocessing by different algorithms has a significant impact. RegRCPqn (regional regression on correlated probes with quantile normalization) does not require raw idat files and can be applied to datasets with only β-values or M-values available, which is a common scenario from real life [44]. RCP [109] is a within-array normalization that uses the spatial correlation of DNA methylation at CpG sites to estimate the calibration transformation between type I and type II intensities. The regRCPqn procedure improves the RCP algorithm by including three functions to solve the problem under study. First, it calculates the RCP normalization separately for each type of genomic region (i.e., for CpG belonging to islands, shores, shelves, or open seas) because the distribution of DNA methylation values is different in each of these types of regions [110]. It then performs a quantile normalization between samples, in which CpG values for all samples are normalized separately for each CpG region and for type I and type II probes. Finally, it introduces the possibility of storing the reference distribution and using it to perform quantile normalization of samples from another dataset based on the reference. The reference distribution is calculated separately for each area type and for type I and type II probes. When possible, this distribution is used by regRCPqn to perform normalization based on the reference, again separately for each region and probe type. The dataset with the highest number of participants for each disease is used as the reference, and the others are harmonized relative to it. We consider CpG sites whose distribution of methylation levels does not differ in different datasets in the control group. To find them, we performed the Mann-Whitney U-test [82] for all CpG sites included in the train datasets for the control group and took CpG sites for which the p-value adjusted according to the Benjamini-Hochberg procedure [111] >0.05. The Mann-Whitney U-test was performed using the scipy package version 1.8.0.

### 4.3. Classification models

The most common type of data for machine learning and deep learning tasks is tabular data, which comprises samples (rows) with the same set of features (columns). DNA methylation is an example of this type of data. Tabular data, unlike image or speech data, is heterogeneous, resulting in dense numerical and sparse categorical features. In addition, the correlation between features is weaker than the spatial or semantic relationship in image or speech data [112]. Variables can be correlated or independent, and features have no positional information. Consequently, it is necessary to detect and use correlation without relying on spatial information [86, 87]. During the last decade, traditional machine learning methods such as gradient-boosted decision trees (GBDT) [83] have continued to dominate tabular data modeling and have demonstrated better performance than deep learning [101]. GBDT trains a series of weak learners to predict the outcome. In GBDT, the weak learner is a standard decision tree that lacks differentiability. Despite their differences, their performance on many problems is similar [84]. When deep neural networks are applied to tabular data, many problems arise, such as lack of locality, missing values, mixed object types (numeric, ordinal and categorical), lack of prior knowledge about the structure of the data. Tree ensemble algorithms are considered a recommended option for real-world problems with tabular data [83, 84, 113]. The XGBoost algorithm [83] is an extendible gradient boosting tree algorithm that achieves state-of-the-art results on many tabular datasets [114, 115].

We consider the following classification models: XGBoost [83], CatBoost [84], LightGBM [85], TabNet [86], NODE [87]. XGBoost (Extreme Gradient Boosting) is a scalable, distributed gradient-boosted decision tree (GBDT) machine learning library. GBDT iteratively trains an ensemble of shallow decision trees, with each iteration using error residuals from the previous model to fit the next model. The final prediction is a weighted sum of all tree predictions. XGBoost has one of the best combinations of prediction performance and processing time. CatBoost is an open-source gradient boosting algorithm, which builds symmetric (balanced) trees. At each step, the leaves of the previous tree are separated by the same condition. A feature-split pair is selected and used for all nodes, which provides the least losses. This balanced tree architecture reduces prediction time and controls overfitting. LightGBM is a fast, distributed, high-performance gradient boosting platform that supports the decision tree algorithm. It splits the tree by leaf with the simplest fit, whereas other boosting algorithms split the tree by depth or by level rather than by leaf. Thus, when growing on an equivalent leaf in LightGBM, a leaf-based algorithm can reduce more losses than a level-based algorithm, and therefore lead to greater accuracy. TabNet is a deep neural network designed to handle tabular data. TabNet inputs raw tabular data with no preprocessing and is trained using gradient descent-based optimization. It uses sequential attention to select features at each decision step, providing interpretability and better learning as the learning capability is used for the most useful features, with instance-specific feature selection. Neural Oblivious Decision Ensembles (NODE) is a deep learning architecture designed to handle tabular data. The NODE architecture generalizes ensembles of oblivious decision trees, but benefits from both end-to-end gradient-based optimization and multilevel hierarchical learning capabilities. All of the above models can handle continuous variables (without categorical ones). For classification, we use only the DNA methylation levels of the different CpG sites, which are continuous variables. Parameter values of the trained models which were found by hyperparametric search can be found in Supplementary Table S3. All models have been trained for 2,000 epochs.

Each model was trained on two training datasets and then tested on the remaining independent datasets. Hyperparametric search was used to find optimal parameters for the models. There are a lot of quality metrics for the classification problem: accuracy, precision, recall, f1 score, Cohen’s kappa, Matthews correlation coefficient, AUROC, etc. As the main metric, we choose accuracy with weighted averaging to take into account the possible imbalance of the classes. Adam optimizer and StepLR scheduler were used for the neural network models [116]. Used versions of software packages for the models: XGBoost 1.5.2, CatBoost 1.0.4, LightGBM 3.3.2, TabNet 3.1.1, PyTorch 1.10.0, PyTorch Lightning 1.6.0.

### 4.4. Dimensionality reduction

Based on the features ranking, we performed dimensionality reduction of the models. We performed it, leaving only the most important CpGs for solving the classification problem. For this purpose, we built a series of models with different numbers of features. First, for each disease, we choose the top 10 most important features, and a new model is built for them (the type of model is chosen beforehand - it is the best in terms of accuracy for the full data). Then new models are built on the number of features from 10 to 1000 in increments of 10, and for each such model, the weighted classification accuracy is calculated. For all these models hyperparametric search was performed. According to the dependence of weighted classification accuracy on the number of features we chose as optimal the number of features for which the highest weighted classification accuracy is observed for the considered diseases.

### 4.5. Imputation of missing values

Missing data can be divided into three classes [117]: i) missing completely at random (MCAR) values, if the probability of absence is completely independent of both observed and unobserved variables; ii) missing at random (MAR) values, if the probability of absence is independent of the value itself, but may depend on observed variables; iii) missing not at random (MNAR), if the probability of absence depends on the missing value itself. There is currently no statistical way to determine which category the specific missing data falls into. Assumptions are usually made based on knowledge of the data and the data collection and processing procedure. It is assumed that the missing values represent MCAR/MAR due to random experimental and technology-related errors [48]. It has been shown that missing values lying at the midrange methylation level are more difficult to impute than missing values close to the extremes of the range [49]. This is probably a consequence of the higher variance of methylation values in the middle ranges. Such a scenario could have a profound effect in terms of performance expectations, assuming that many missing values in the data are of the MNAR type and, in particular, lie in the middle range of β values.

In general terms, imputation approaches can be divided into single (SI) and multiple imputation (MI) methods. SI methods replace a missing value with a single acceptable value. MI methods perform multiple SIs and average parameter estimates over multiple imputations to produce a single estimate. Under MCAR/MAR assumptions, the most common imputation methods like mean, median or mode can handle missing data [118]. Such simple imputation methods are used often [119], but they can lead to systematic error or unrealistic results for multivariate datasets. In addition, for large data, this method often performs poorly [120]. The expectation maximization method is an iterative method for handling missing values in numerical datasets, and the algorithm uses an “impute, estimate, and iterate until convergence” approach. Each iteration involves two steps: expectation and maximization. Expectation estimates the missing values given the observed data, while maximization uses the current estimated values to maximize the probability of all data [121–123]. Besides classical methods, there are approaches to multiple imputation, for example, chained equation for big data [120]. Hot-deck imputation handles missing values by matching missing values with other values in the dataset for several other key variables that have complete values [124, 125]. However, this method does not account for the variability of the missing data. One of the common hot-deck methods is K Nearest Neighbours (KNN) [126]. The KNN algorithm works by classifying the nearest neighbors of missing values and using those neighbors for imputation using a distance measure between instances [127]. Several distance measures can be used for KNN imputation, but Euclidean distance has been shown to provide efficiency and performance [128] and is therefore the most widely used distance measure. However, KNN imputation has weaknesses, such as poor accuracy when imputing variables and introducing false associations where none exists [129]. Another weakness of KNN imputation is that it scans the entire dataset, which increases computation time [130]. However, there are approaches developed in the literature to improve the KNN imputation algorithm [131–137]. All imputation methods that can deal with continuous variables are suitable for imputing DNA methylation data [48]. To study the effect of these methods on classification accuracy, we removed from consideration 100 CpG sites with the highest importance values for each disease and tried to fill them in. We used previously constructed small models for both diseases. Imputation methods were applied by impyute package version 0.0.8.

### 4.6. Explainable artificial intelligence

Modern machine-learning-based artificial intelligence systems are usually treated as black boxes. However, every decision must be made available for verification by a human expert [138]. One important aspect of model explainability is the ability to verify the system. For example, in healthcare, the use of models that can be interpreted and verified by medical experts is an absolute necessity [139]. Another aspect is to improve the system. The first step to improving the AI system is to understand its weaknesses. Performing weakness analysis on black box models is more difficult than on models that can be interpreted. Furthermore, model interpretability can be useful when comparing different models or architectures [140–142]. It can be argued that the better we understand what models do (and why they sometimes fail), the easier it becomes to improve them [138]. The next important aspect of explainability is the ability to learn from the system: since modern AI systems learn from millions of examples, they can observe patterns in the data that are inaccessible to humans, who can only learn from a limited number of examples [143, 144]. Explainability is also important for other machine learning methods beyond neural networks [140]. One of the taxonomies to classify explanatory methods is global and local methods [58, 59, 138]. Local interpretable methods apply to a single model result; they can explain the reason for a particular prediction or result. In contrast, global methods try to explain the behavior of the model as a whole. Perturbation is the easiest way to analyze the effect of changing input features on the AI model outputs. This can be accomplished by removing or changing certain input features, running a forward pass, and measuring the difference with the original output data. The input characteristics that most affect the output are evaluated as the most important ones. This is computationally costly, since a direct pass must be run after perturbing each group of input features. Such a perturbation-based approach is Shapley value sampling, which computes approximate Shapley values by taking each input feature for a certain number of times. It is a method from game theory that describes a fair distribution of wins and losses between input functions [145]. As a result, it is not a practical method in its original form, but has led to the development of methods based on game theory, such as Deep SHapley Additive exPlanations (SHAP) [97]. SHAP has an alternative kernel-based approach to estimating Shapley values inspired by local surrogate models. There is also TreeSHAP, an efficient approach to estimating tree models, as well as DeepExplainer, an enhanced version of the DeepLIFT algorithm for deep neural networks. For the constructed portable models, we applied SHAP to obtain global and local explainability. SHAP values were calculated using eponymous package version 0.40.0.

## Data availability statement

No new data was generated. Data used in this study are available from the GEO database (accession numbers GSE145361, GSE111629, GSE72774, GSE84727, GSE80417, GSE152027, GSE116379).

## Code availability statement

The source code for the analysis pipeline presented in the manuscript is publicly available.

Project name: DNAmClassMeta

Project home page: https://github.com/GillianGrayson/DNAmClassMeta

Operating system(s): Platform independent

Programming language: Python

Other requirements: Python 3.8 or higher, pytorch-lightning 1.5.10 or higher, xgboost 1.6.0 or higher, catboost 1.0.5 or higher, lightgbm 3.3.2 or higher. All requirements are listed in the requirements.txt file in the project home page.

License: MIT

## Abbreviations

AI: Artificial Intelligence; CatBoost: Categorical Boosting; DMP: Differentially Methylated Position; DMR: Differentially Methylated Region; DNAm: DNA methylation; EWAS: Epigenome-Wide Association Study; FDR: False Discovery Rate; GBDT: Gradient-Boosted Decision Tree; IPD: Individual Participant Data; KNN: K Nearest Neighbors; LightGBM: Light Gradient Boosting Machine; MAR: Missing At Random; MCAR: Missing Completely At Random; MI: Multiple Imputation; MNAR: Missing Not At Random; NODE: Neural Oblivious Decision Ensemble; PTSD: Post-Traumatic Stress Disorder; RCP: Regression on Correlated Probes; SHAP: Shapley Additive Explanations; SI: Single Imputation; XAI: Explainable Artificial Intelligence; XGBoost: Extreme Gradient Boosting.

## Author’s Contributions

Conceptualization: A.K., I.Y., M.G.B., M.I.; Formal analysis: A.K., I.Y.; Methodology: A.K., I.Y., M.G.B.; Software: A.K., I.Y.; Supervision: M.G.B., C.F., M.V., M.I.; Visualization: A.K., I.Y.; Writing – original draft: A.K., I.Y.; Writing – review & editing: A.K., I.Y., M.G.B., C.F., M.V., M.I.

## Acknowledgements

The authors acknowledge the use of computational resources provided by the “Lobachevsky” supercomputer.

## References

1. Sasaki H, Matsui Y (2008) Epigenetic events in mammalian germ-cell development: reprogramming and beyond. Nat Rev Genet 9:129–140. https://doi.org/10.1038/nrg2295

2. Igarashi J, Muroi S, Kawashima H, Wang X, Shinojima Y, Kitamura E, Oinuma T, Nemoto N, Song F, Ghosh S, Held WA, Nagase H (2008) Quantitative analysis of human tissue-specific differences in methylation. Biochem Biophys Res Commun 376:658–664. https://doi.org/10.1016/j.bbrc.2008.09.044

3. Zemach A, McDaniel IE, Silva P, Zilberman D (2010) Genome-wide evolutionary analysis of eukaryotic DNA methylation. Science 328:916–919. https://doi.org/10.1126/science.1186366

4. Ziller MJ, Gu H, Müller F, Donaghey J, Tsai LT-Y, Kohlbacher O, De Jager PL, Rosen ED, Bennett DA, Bernstein BE, Gnirke A, Meissner A (2013) Charting a dynamic DNA methylation landscape of the human genome. Nature 500:477–481. https://doi.org/10.1038/nature12433

5. Horvath S (2013) DNA methylation age of human tissues and cell types. Genome Biol 14:R115. https://doi.org/10.1186/gb-2013-14-10-r115

6. Orozco LD, Farrell C, Hale C, Rubbi L, Rinaldi A, Civelek M, Pan C, Lam L, Montoya D, Edillor C, Seldin M, Boehnke M, Mohlke KL, Jacobsen S, Kuusisto J, Laakso M, Lusis AJ, Pellegrini M (2018) Epigenome-wide association in adipose tissue from the METSIM cohort. Hum Mol Genet 27:2586. https://doi.org/10.1093/hmg/ddy205

7. Smith ZD, Meissner A (2013) DNA methylation: roles in mammalian development. Nat Rev Genet 14:204–220. https://doi.org/10.1038/nrg3354

8. Lim DHK, Maher ER (2010) Genomic imprinting syndromes and cancer. Adv Genet 70:145–175. https://doi.org/10.1016/B978-0-12-380866-0.60006-X

9. Robertson KD (2005) DNA methylation and human disease. Nat Rev Genet 6:597–610. https://doi.org/10.1038/nrg1655

10. Christensen BC, Houseman EA, Marsit CJ, Zheng S, Wrensch MR, Wiemels JL, Nelson HH, Karagas MR, Padbury JF, Bueno R, Sugarbaker DJ, Yeh R-F, Wiencke JK, Kelsey KT (2009) Aging and environmental exposures alter tissue-specific DNA methylation dependent upon CpG island context. PLoS Genet 5:e1000602. https://doi.org/10.1371/journal.pgen.1000602

11. Bell CG, Lowe R, Adams PD, Baccarelli AA, Beck S, Bell JT, Christensen BC, Gladyshev VN, Heijmans BT, Horvath S, Ideker T, Issa J-PJ, Kelsey KT, Marioni RE, Reik W, Relton CL, Schalkwyk LC, Teschendorff AE, Wagner W, Zhang K, Rakyan VK (2019) DNA methylation aging clocks: challenges and recommendations. Genome Biol 20:249. https://doi.org/10.1186/s13059-019-1824-y

12. Moran S, Arribas C, Esteller M (2016) Validation of a DNA methylation microarray for 850,000 CpG sites of the human genome enriched in enhancer sequences. Epigenomics 8:389–399. https://doi.org/10.2217/epi.15.114

13. Bibikova M, Lin Z, Zhou L, Chudin E, Garcia EW, Wu B, Doucet D, Thomas NJ, Wang Y, Vollmer E, Goldmann T, Seifart C, Jiang W, Barker DL, Chee MS, Floros J, Fan J-B (2006) High-throughput DNA methylation profiling using universal bead arrays. Genome Res 16:383–393. https://doi.org/10.1101/gr.4410706

14. Irizarry RA, Ladd-Acosta C, Carvalho B, Wu H, Brandenburg SA, Jeddeloh JA, Wen B, Feinberg AP (2008) Comprehensive high-throughput arrays for relative methylation (CHARM). Genome Res 18:780–790. https://doi.org/10.1101/gr.7301508

15. Du P, Zhang X, Huang C-C, Jafari N, Kibbe WA, Hou L, Lin SM (2010) Comparison of Beta-value and M-value methods for quantifying methylation levels by microarray analysis. BMC Bioinformatics 11:587. https://doi.org/10.1186/1471-2105-11-587

16. Tian T, Wan J, Song Q, Wei Z (2019) Clustering single-cell RNA-seq data with a model-based deep learning approach. Nat Mach Intell 1:191–198. https://doi.org/10.1038/s42256-019-0037-0

17. Lopez R, Regier J, Cole MB, Jordan MI, Yosef N (2018) Deep generative modeling for single-cell transcriptomics. Nat Methods 15:1053–1058. https://doi.org/10.1038/s41592-018-0229-2

18. Way GP, Greene CS (2018) Extracting a biologically relevant latent space from cancer transcriptomes with variational autoencoders. Pac Symp Biocomput 23:80–91

19. Titus AJ, Wilkins OM, Bobak CA, Christensen BC (2018) Unsupervised deep learning with variational autoencoders applied to breast tumor genome-wide DNA methylation data with biologic feature extraction. Bioinformatics

20. Ching T, Himmelstein DS, Beaulieu-Jones BK, Kalinin AA, Do BT, Way GP, Ferrero E, Agapow P-M, Zietz M, Hoffman MM, Xie W, Rosen GL, Lengerich BJ, Israeli J, Lanchantin J, Woloszynek S, Carpenter AE, Shrikumar A, Xu J, Cofer EM, Lavender CA, Turaga SC, Alexandari AM, Lu Z, Harris DJ, DeCaprio D, Qi Y, Kundaje A, Peng Y, Wiley LK, Segler MHS, Boca SM, Swamidass SJ, Huang A, Gitter A, Greene CS (2018) Opportunities and obstacles for deep learning in biology and medicine. J R Soc Interface 15:20170387. https://doi.org/10.1098/rsif.2017.0387

21. Levy JJ, Titus AJ, Petersen CL, Chen Y, Salas LA, Christensen BC (2020) MethylNet: an automated and modular deep learning approach for DNA methylation analysis. BMC Bioinformatics 21:108. https://doi.org/10.1186/s12859-020-3443-8

22. The Cancer Genome Atlas Research Network, Weinstein JN, Collisson EA, Mills GB, Shaw KRM, Ozenberger BA, Ellrott K, Shmulevich I, Sander C, Stuart JM (2013) The Cancer Genome Atlas Pan-Cancer analysis project. Nat Genet 45:1113–1120. https://doi.org/10.1038/ng.2764

23. Ding W, Chen G, Shi T (2019) Integrative analysis identifies potential DNA methylation biomarkers for pan-cancer diagnosis and prognosis. Epigenetics 14:67–80. https://doi.org/10.1080/15592294.2019.1568178

24. Celli F, Cumbo F, Weitschek E (2018) Classification of Large DNA Methylation Datasets for Identifying Cancer Drivers. Big Data Research 13:21–28. https://doi.org/10.1016/j.bdr.2018.02.005

25. Ma B, Meng F, Yan G, Yan H, Chai B, Song F (2020) Diagnostic classification of cancers using extreme gradient boosting algorithm and multi-omics data. Computers in Biology and Medicine 121:103761. https://doi.org/10.1016/j.compbiomed.2020.103761

26. List M, Hauschild A-C, Tan Q, Kruse TA, Baumbach J, Batra R (2014) Classification of Breast Cancer Subtypes by combining Gene Expression and DNA Methylation Data. Journal of Integrative Bioinformatics 11:1–14. https://doi.org/10.1515/jib-2014-236

27. Dong R, Yang X, Zhang X, Gao P, Ke A, Sun H, Zhou J, Fan J, Cai J, Shi G (2019) Predicting overall survival of patients with hepatocellular carcinoma using a three-category method based on DNA methylation and machine learning. J Cell Mol Med 23:3369–3374. https://doi.org/10.1111/jcmm.14231

28. Hao X, Luo H, Krawczyk M, Wei W, Wang W, Wang J, Flagg K, Hou J, Zhang H, Yi S, Jafari M, Lin D, Chung C, Caughey BA, Li G, Dhar D, Shi W, Zheng L, Hou R, Zhu J, Zhao L, Fu X, Zhang E, Zhang C, Zhu J-K, Karin M, Xu R-H, Zhang K (2017) DNA methylation markers for diagnosis and prognosis of common cancers. Proc Natl Acad Sci USA 114:7414–7419. https://doi.org/10.1073/pnas.1703577114

29. Jurmeister P, Bockmayr M, Seegerer P, Bockmayr T, Treue D, Montavon G, Vollbrecht C, Arnold A, Teichmann D, Bressem K, Schüller U, von Laffert M, Müller K-R, Capper D, Klauschen F (2019) Machine learning analysis of DNA methylation profiles distinguishes primary lung squamous cell carcinomas from head and neck metastases. Sci Transl Med 11:eaaw8513. https://doi.org/10.1126/scitranslmed.aaw8513

30. Wajed SA, Laird PW, DeMeester TR (2001) DNA Methylation: An Alternative Pathway to Cancer: Annals of Surgery 234:10–20. https://doi.org/10.1097/00000658-200107000-00003

31. Bollepalli S, Korhonen T, Kaprio J, Anders S, Ollikainen M (2019) EpiSmokEr: a robust classifier to determine smoking status from DNA methylation data. Epigenomics 11:1469–1486. https://doi.org/10.2217/epi-2019-0206

32. Lee Y-C, Christensen JJ, Parnell LD, Smith CE, Shao J, McKeown NM, Ordovás JM, Lai C-Q (2022) Using Machine Learning to Predict Obesity Based on Genome-Wide and Epigenome-Wide Gene–Gene and Gene–Diet Interactions. Front Genet 12:783845. https://doi.org/10.3389/fgene.2021.783845

33. Aref-Eshghi E, Rodenhiser DI, Schenkel LC, Lin H, Skinner C, Ainsworth P, Paré G, Hood RL, Bulman DE, Kernohan KD, Care4Rare Canada Consortium, Boycott KM, Campeau PM, Schwartz C, Sadikovic B (2018) Genomic DNA Methylation Signatures Enable Concurrent Diagnosis and Clinical Genetic Variant Classification in Neurodevelopmental Syndromes. Am J Hum Genet 102:156–174. https://doi.org/10.1016/j.ajhg.2017.12.008

34. Dogan MV, Grumbach IM, Michaelson JJ, Philibert RA (2018) Integrated genetic and epigenetic prediction of coronary heart disease in the Framingham Heart Study. PLoS One 13:e0190549. https://doi.org/10.1371/journal.pone.0190549

35. Gunasekara CJ, Hannon E, MacKay H, Coarfa C, McQuillin A, Clair DSt, Mill J, Waterland RA (2021) A machine learning case–control classifier for schizophrenia based on DNA methylation in blood. Transl Psychiatry 11:412. https://doi.org/10.1038/s41398-021-01496-3

36. Jabari S, Kobow K, Pieper T, Hartlieb T, Kudernatsch M, Polster T, Bien CG, Kalbhenn T, Simon M, Hamer H, Rössler K, Feucht M, Mühlebner A, Najm I, Peixoto-Santos JE, Gil-Nagel A, Delgado RT, Aledo-Serrano A, Hou Y, Coras R, von Deimling A, Blümcke I (2022) DNA methylation-based classification of malformations of cortical development in the human brain. Acta Neuropathol 143:93–104. https://doi.org/10.1007/s00401-021-02386-0

37. Jo T, Nho K, Bice P, Saykin AJ, for the Alzheimer’s Neuroimaging Initiative (2021) Deep learning-based identification of genetic variants: Application to Alzheimer’s disease classification. Genetic and Genomic Medicine

38. Haghshenas S, Bhai P, Aref-Eshghi E, Sadikovic B (2020) Diagnostic Utility of Genome-Wide DNA Methylation Analysis in Mendelian Neurodevelopmental Disorders. IJMS 21:9303. https://doi.org/10.3390/ijms21239303

39. Xiong Z, Zhang X, Zhang M, Cao B (2020) Predicting Features of Human Mental Disorders through Methylation Profile and Machine Learning Models. In: 2020 2nd International Conference on Machine Learning, Big Data and Business Intelligence (MLBDBI). IEEE, Taiyuan, China, pp 67–75

40. Luo X, Wei Y (2019) Batch Effects Correction with Unknown Subtypes. Journal of the American Statistical Association 114:581–594. https://doi.org/10.1080/01621459.2018.1497494

41. Leek JT, Scharpf RB, Bravo HC, Simcha D, Langmead B, Johnson WE, Geman D, Baggerly K, Irizarry RA (2010) Tackling the widespread and critical impact of batch effects in high-throughput data. Nat Rev Genet 11:733–739. https://doi.org/10.1038/nrg2825

42. Perrier F, Novoloaca A, Ambatipudi S, Baglietto L, Ghantous A, Perduca V, Barrdahl M, Harlid S, Ong KK, Cardona A, Polidoro S, Nøst TH, Overvad K, Omichessan H, Dollé M, Bamia C, Huerta JM, Vineis P, Herceg Z, Romieu I, Ferrari P (2018) Identifying and correcting epigenetics measurements for systematic sources of variation. Clin Epigenet 10:38. https://doi.org/10.1186/s13148-018-0471-6

43. Zindler T, Frieling H, Neyazi A, Bleich S, Friedel E (2020) Simulating ComBat: how batch correction can lead to the systematic introduction of false positive results in DNA methylation microarray studies. BMC Bioinformatics 21:271. https://doi.org/10.1186/s12859-020-03559-6

44. Sala C, Di Lena P, Fernandes Durso D, Prodi A, Castellani G, Nardini C (2020) Evaluation of pre-processing on the meta-analysis of DNA methylation data from the Illumina HumanMethylation450 BeadChip platform. PLoS One 15:e0229763. https://doi.org/10.1371/journal.pone.0229763

45. Garagnani P, Bacalini MG, Pirazzini C, Gori D, Giuliani C, Mari D, Di Blasio AM, Gentilini D, Vitale G, Collino S, Rezzi S, Castellani G, Capri M, Salvioli S, Franceschi C (2012) Methylation of ELOVL2 gene as a new epigenetic marker of age. Aging Cell 11:1132–1134. https://doi.org/10.1111/acel.12005

46. Hannum G, Guinney J, Zhao L, Zhang L, Hughes G, Sadda S, Klotzle B, Bibikova M, Fan J-B, Gao Y, Deconde R, Chen M, Rajapakse I, Friend S, Ideker T, Zhang K (2013) Genome-wide Methylation Profiles Reveal Quantitative Views of Human Aging Rates. Molecular Cell 49:359–367. https://doi.org/10.1016/j.molcel.2012.10.016

47. Weidner C, Lin Q, Koch C, Eisele L, Beier F, Ziegler P, Bauerschlag D, Jöckel K-H, Erbel R, Mühleisen T, Zenke M, Brümmendorf T, Wagner W (2014) Aging of blood can be tracked by DNA methylation changes at just three CpG sites. Genome Biol 15:R24. https://doi.org/10.1186/gb-2014-15-2-r24

48. Di Lena P, Sala C, Prodi A, Nardini C (2019) Missing value estimation methods for DNA methylation data. Bioinformatics 35:3786–3793. https://doi.org/10.1093/bioinformatics/btz134

49. Lena PD, Sala C, Prodi A, Nardini C (2020) Methylation data imputation performances under different representations and missingness patterns. BMC Bioinformatics 21:268. https://doi.org/10.1186/s12859-020-03592-5

50. Venkat N (2018) The Curse of Dimensionality: Inside Out. https://doi.org/10.13140/RG.2.2.29631.36006

51. Amor R del, Colomer A, Monteagudo C, Naranjo V (2021) A deep embedded refined clustering approach for breast cancer distinction based on DNA methylation. Neural Comput & Applic. https://doi.org/10.1007/s00521-021-06357-0

52. Levine ME, Lu AT, Quach A, Chen BH, Assimes TL, Bandinelli S, Hou L, Baccarelli AA, Stewart JD, Li Y, Whitsel EA, Wilson JG, Reiner AP, Aviv A, Lohman K, Liu Y, Ferrucci L, Horvath S (2018) An epigenetic biomarker of aging for lifespan and healthspan. Aging 10:573–591. https://doi.org/10.18632/aging.101414

53. Lu AT, Quach A, Wilson JG, Reiner AP, Aviv A, Raj K, Hou L, Baccarelli AA, Li Y, Stewart JD, Whitsel EA, Assimes TL, Ferrucci L, Horvath S (2019) DNA methylation GrimAge strongly predicts lifespan and healthspan. Aging 11:303–327. https://doi.org/10.18632/aging.101684

54. Kurdyukov S, Bullock M (2016) DNA Methylation Analysis: Choosing the Right Method. Biology (Basel) 5:E3. https://doi.org/10.3390/biology5010003

55. He K, Zhang X, Ren S, Sun J (2016) Deep Residual Learning for Image Recognition. In: 2016 IEEE Conference on Computer Vision and Pattern Recognition (CVPR). IEEE, Las Vegas, NV, USA, pp 770–778

56. Cho K, van Merriënboer B, Gulcehre C, Bahdanau D, Bougares F, Schwenk H, Bengio Y (2014) Learning Phrase Representations using RNN Encoder–Decoder for Statistical Machine Translation. In: Proceedings of the 2014 Conference on Empirical Methods in Natural Language Processing (EMNLP). Association for Computational Linguistics, Doha, Qatar, pp 1724–1734

57. Deng L, Hinton G, Kingsbury B (2013) New types of deep neural network learning for speech recognition and related applications: an overview. In: 2013 IEEE International Conference on Acoustics, Speech and Signal Processing. pp 8599–8603

58. Baehrens D, Schroeter T, Harmeling S, Kawanabe M, Hansen K, Müller K-R (2010) How to Explain Individual Classification Decisions. J Mach Learn Res 11:1803–1831

59. Simonyan K, Vedaldi A, Zisserman A (2014) Deep Inside Convolutional Networks: Visualising Image Classification Models and Saliency Maps. arXiv:13126034 [cs]

60. Zeiler MD, Fergus R (2014) Visualizing and Understanding Convolutional Networks. In: Fleet D, Pajdla T, Schiele B, Tuytelaars T (eds) Computer Vision – ECCV 2014. Springer International Publishing, Cham, pp 818–833

61. Bach S, Binder A, Montavon G, Klauschen F, Müller K-R, Samek W (2015) On Pixel-Wise Explanations for Non-Linear Classifier Decisions by Layer-Wise Relevance Propagation. PLoS ONE 10:e0130140. https://doi.org/10.1371/journal.pone.0130140

62. Shrikumar A, Greenside P, Shcherbina A, Kundaje A (2017) Not Just a Black Box: Learning Important Features Through Propagating Activation Differences. arXiv:160501713 [cs]

63. Mahendran A, Vedaldi A (2016) Visualizing Deep Convolutional Neural Networks Using Natural Pre-images. Int J Comput Vis 120:233–255. https://doi.org/10.1007/s11263-016-0911-8

64. Lipton ZC (2017) The Mythos of Model Interpretability. arXiv:160603490 [cs, stat]

65. Ribeiro MT, Singh S, Guestrin C (2016) “Why Should I Trust You?”: Explaining the Predictions of Any Classifier. arXiv:160204938 [cs, stat]

66. Zintgraf LM, Cohen TS, Adel T, Welling M (2017) Visualizing Deep Neural Network Decisions: Prediction Difference Analysis. arXiv:170204595 [cs]

67. Doshi-Velez F, Kim B (2017) Towards A Rigorous Science of Interpretable Machine Learning. arXiv:170208608 [cs, stat]

68. Montavon G, Samek W, Müller K-R (2018) Methods for interpreting and understanding deep neural networks. Digital Signal Processing 73:1–15. https://doi.org/10.1016/j.dsp.2017.10.011

69. Mahendran A, Vedaldi A (2014) Understanding Deep Image Representations by Inverting Them. arXiv:14120035 [cs]

70. Nguyen A, Yosinski J, Clune J (2016) Multifaceted Feature Visualization: Uncovering the Different Types of Features Learned By Each Neuron in Deep Neural Networks. arXiv:160203616 [cs]

71. Landecker W, Thomure MD, Bettencourt LMA, Mitchell M, Kenyon GT, Brumby SP (2013) Interpreting individual classifications of hierarchical networks. In: 2013 IEEE Symposium on Computational Intelligence and Data Mining (CIDM). pp 32–38

72. Montavon G, Lapuschkin S, Binder A, Samek W, Müller K-R (2017) Explaining nonlinear classification decisions with deep Taylor decomposition. Pattern Recogn 65:211–222. https://doi.org/10.1016/j.patcog.2016.11.008

73. Vallerga CL, Zhang F, Fowdar J, McRae AF, Qi T, Nabais MF, Zhang Q, Kassam I, Henders AK, Wallace L, Montgomery G, Chuang Y-H, Horvath S, Ritz B, Halliday G, Hickie I, Kwok JB, Pearson J, Pitcher T, Kennedy M, Bentley SR, Silburn PA, Yang J, Wray NR, Lewis SJG, Anderson T, Dalrymple-Alford J, Mellick GD, Visscher PM, Gratten J (2020) Analysis of DNA methylation associates the cystine-glutamate antiporter SLC7A11 with risk of Parkinson’s disease. Nat Commun 11:1238. https://doi.org/10.1038/s41467-020-15065-7

74. Chuang Y-H, Paul KC, Bronstein JM, Bordelon Y, Horvath S, Ritz B (2017) Parkinson’s disease is associated with DNA methylation levels in human blood and saliva. Genome Med 9:76. https://doi.org/10.1186/s13073-017-0466-5

75. Horvath S, Ritz BR (2015) Increased epigenetic age and granulocyte counts in the blood of Parkinson’s disease patients. Aging (Albany NY) 7:1130–1142. https://doi.org/10.18632/aging.100859

76. Chuang Y-H, Lu AT, Paul KC, Folle AD, Bronstein JM, Bordelon Y, Horvath S, Ritz B (2019) Longitudinal Epigenome-Wide Methylation Study of Cognitive Decline and Motor Progression in Parkinson’s Disease. J Parkinsons Dis 9:389–400. https://doi.org/10.3233/JPD-181549

77. Paul KC, Binder AM, Horvath S, Kusters C, Yan Q, Rosario ID, Yu Y, Bronstein J, Ritz B (2021) Accelerated hematopoietic mitotic aging measured by DNA methylation, blood cell lineage, and Parkinson’s disease. BMC Genomics 22:696. https://doi.org/10.1186/s12864-021-08009-y

78. Hannon E, Dempster EL, Mansell G, Burrage J, Bass N, Bohlken MM, Corvin A, Curtis CJ, Dempster D, Di Forti M, Dinan TG, Donohoe G, Gaughran F, Gill M, Gillespie A, Gunasinghe C, Hulshoff HE, Hultman CM, Johansson V, Kahn RS, Kaprio J, Kenis G, Kowalec K, MacCabe J, McDonald C, McQuillin A, Morris DW, Murphy KC, Mustard CJ, Nenadic I, O’Donovan MC, Quattrone D, Richards AL, Rutten BP, St Clair D, Therman S, Toulopoulou T, Van Os J, Waddington JL, Wellcome Trust Case Control Consortium (WTCCC), CRESTAR consortium, Sullivan P, Vassos E, Breen G, Collier DA, Murray RM, Schalkwyk LS, Mill J (2021) DNA methylation meta-analysis reveals cellular alterations in psychosis and markers of treatment-resistant schizophrenia. Elife 10:e58430. https://doi.org/10.7554/eLife.58430

79. Hannon E, Dempster E, Viana J, Burrage J, Smith AR, Macdonald R, St Clair D, Mustard C, Breen G, Therman S, Kaprio J, Toulopoulou T, Hulshoff Pol HE, Bohlken MM, Kahn RS, Nenadic I, Hultman CM, Murray RM, Collier DA, Bass N, Gurling H, McQuillin A, Schalkwyk L, Mill J (2016) An integrated genetic-epigenetic analysis of schizophrenia: evidence for co-localization of genetic associations and differential DNA methylation. Genome Biol 17:176. https://doi.org/10.1186/s13059-016-1041-x

80. Boks MP, Houtepen LC, Xu Z, He Y, Ursini G, Maihofer AX, Rajarajan P, Yu Q, Xu H, Wu Y, Wang S, Shi JP, Hulshoff Pol HE, Strengman E, Rutten BPF, Jaffe AE, Kleinman JE, Baker DG, Hol EM, Akbarian S, Nievergelt CM, De Witte LD, Vinkers CH, Weinberger DR, Yu J, Kahn RS (2018) Genetic vulnerability to DUSP22 promoter hypermethylation is involved in the relation between in utero famine exposure and schizophrenia. NPJ Schizophr 4:16. https://doi.org/10.1038/s41537-018-0058-4

81. Rauschert S, Raubenheimer K, Melton PE, Huang RC (2020) Machine learning and clinical epigenetics: a review of challenges for diagnosis and classification. Clin Epigenet 12:51. https://doi.org/10.1186/s13148-020-00842-4

82. Mann HB, Whitney DR (1947) On a Test of Whether one of Two Random Variables is Stochastically Larger than the Other. Ann Math Statist 18:50–60. https://doi.org/10.1214/aoms/1177730491

83. Chen T, Guestrin C (2016) XGBoost: A Scalable Tree Boosting System. In: Proceedings of the 22nd ACM SIGKDD International Conference on Knowledge Discovery and Data Mining. ACM, San Francisco California USA, pp 785–794

84. Prokhorenkova L, Gusev G, Vorobev A, Dorogush AV, Gulin A (2018) CatBoost: unbiased boosting with categorical features. In: Bengio S, Wallach H, Larochelle H, Grauman K, Cesa-Bianchi N, Garnett R (eds) Advances in Neural Information Processing Systems. Curran Associates, Inc.

85. Ke G, Meng Q, Finley T, Wang T, Chen W, Ma W, Ye Q, Liu T-Y (2017) LightGBM: A Highly Efficient Gradient Boosting Decision Tree. Long Beach, CA, USA

86. Arik SO, Pfister T (2020) TabNet: Attentive Interpretable Tabular Learning. arXiv:190807442 [cs, stat]

87. Popov S, Morozov S, Babenko A (2019) Neural Oblivious Decision Ensembles for Deep Learning on Tabular Data. arXiv:190906312 [cs, stat]

88. Henderson-Smith A, Fisch KM, Hua J, Liu G, Ricciardelli E, Jepsen K, Huentelman M, Stalberg G, Edland SD, Scherzer CR, Dunckley T, Desplats P (2019) DNA methylation changes associated with Parkinson’s disease progression: outcomes from the first longitudinal genome-wide methylation analysis in blood. Epigenetics 14:365–382. https://doi.org/10.1080/15592294.2019.1588682

89. Kaut O, Schmitt I, Tost J, Busato F, Liu Y, Hofmann P, Witt SH, Rietschel M, Fröhlich H, Wüllner U (2017) Epigenome-wide DNA methylation analysis in siblings and monozygotic twins discordant for sporadic Parkinson’s disease revealed different epigenetic patterns in peripheral blood mononuclear cells. Neurogenetics 18:7–22. https://doi.org/10.1007/s10048-016-0497-x

90. Walton E, Hass J, Liu J, Roffman JL, Bernardoni F, Roessner V, Kirsch M, Schackert G, Calhoun V, Ehrlich S (2016) Correspondence of DNA Methylation Between Blood and Brain Tissue and Its Application to Schizophrenia Research. SCHBUL 42:406–414. https://doi.org/10.1093/schbul/sbv074

91. Hoang HT, Schlager MA, Carter AP, Bullock SL (2017) DYNC1H1 mutations associated with neurological diseases compromise processivity of dynein–dynactin–cargo adaptor complexes. Proc Natl Acad Sci USA 114:. https://doi.org/10.1073/pnas.1620141114

92. Chen X-J, Xu H, Cooper HM, Liu Y (2014) Cytoplasmic dynein: a key player in neurodegenerative and neurodevelopmental diseases. Sci China Life Sci 57:372–377. https://doi.org/10.1007/s11427-014-4639-9

93. Ma Y, Li J, Xu Y, Wang Y, Yao Y, Liu Q, Wang M, Zhao X, Fan R, Chen J, Zhang B, Cai Z, Han H, Yang Z, Yuan W, Zhong Y, Chen X, Ma JZ, Payne TJ, Xu Y, Ning Y, Cui W, Li MD (2020) Identification of 34 genes conferring genetic and pharmacological risk for the comorbidity of schizophrenia and smoking behaviors. Aging (Albany NY) 12:2169–2225. https://doi.org/10.18632/aging.102735

94. Peykov S, Berkel S, Schoen M, Weiss K, Degenhardt F, Strohmaier J, Weiss B, Proepper C, Schratt G, Nöthen MM, Boeckers TM, Rietschel M, Rappold GA (2015) Identification and functional characterization of rare SHANK2 variants in schizophrenia. Mol Psychiatry 20:1489–1498. https://doi.org/10.1038/mp.2014.172

95. Chen X, Long F, Cai B, Chen X, Chen G (2017) A novel relationship for schizophrenia, bipolar and major depressive disorder Part 5: a hint from chromosome 5 high density association screen. Am J Transl Res 9:2473–2491

96. Hindley G, Bahrami S, Steen NE, O’Connell KS, Frei O, Shadrin A, Bettella F, Rødevand L, Fan CC, Dale AM, Djurovic S, Smeland OB, Andreassen OA (2021) Characterising the shared genetic determinants of bipolar disorder, schizophrenia and risk-taking. Transl Psychiatry 11:466. https://doi.org/10.1038/s41398-021-01576-4

97. Chen H, Lundberg S, Lee S-I (2021) Explaining Models by Propagating Shapley Values of Local Components. In: Shaban-Nejad A, Michalowski M, Buckeridge DL (eds) Explainable AI in Healthcare and Medicine: Building a Culture of Transparency and Accountability. Springer International Publishing, Cham, pp 261–270

98. Smyth GK, Speed T (2003) Normalization of cDNA microarray data. Methods 31:265–273. https://doi.org/10.1016/s1046-2023(03)00155-5

99. Johnson WE, Li C, Rabinovic A (2007) Adjusting batch effects in microarray expression data using empirical Bayes methods. Biostatistics 8:118–127. https://doi.org/10.1093/biostatistics/kxj037

100. Price EM, Robinson WP (2018) Adjusting for Batch Effects in DNA Methylation Microarray Data, a Lesson Learned. Front Genet 9:83. https://doi.org/10.3389/fgene.2018.00083

101. Shwartz-Ziv R, Armon A (2022) Tabular data: Deep learning is not all you need. Information Fusion 81:84–90. https://doi.org/10.1016/j.inffus.2021.11.011

102. INTRuST Clinical Consortium, VA Mid-Atlantic MIRECC Workgroup, PGC PTSD Epigenetics Workgroup, Smith AK, Ratanatharathorn A, Maihofer AX, Naviaux RK, Aiello AE, Amstadter AB, Ashley-Koch AE, Baker DG, Beckham JC, Boks MP, Bromet E, Dennis M, Galea S, Garrett ME, Geuze E, Guffanti G, Hauser MA, Katrinli S, Kilaru V, Kessler RC, Kimbrel NA, Koenen KC, Kuan P-F, Li K, Logue MW, Lori A, Luft BJ, Miller MW, Naviaux JC, Nugent NR, Qin X, Ressler KJ, Risbrough VB, Rutten BPF, Stein MB, Ursano RJ, Vermetten E, Vinkers CH, Wang L, Youssef NA, Uddin M, Nievergelt CM (2020) Epigenome-wide meta-analysis of PTSD across 10 military and civilian cohorts identifies methylation changes in AHRR. Nat Commun 11:5965. https://doi.org/10.1038/s41467-020-19615-x

103. Barrett T, Troup DB, Wilhite SE, Ledoux P, Rudnev D, Evangelista C, Kim IF, Soboleva A, Tomashevsky M, Marshall KA, Phillippy KH, Sherman PM, Muertter RN, Edgar R (2009) NCBI GEO: archive for high-throughput functional genomic data. Nucleic Acids Res 37:D885–890. https://doi.org/10.1093/nar/gkn764

104. McCartney DL, Walker RM, Morris SW, McIntosh AM, Porteous DJ, Evans KL (2016) Identification of polymorphic and off-target probe binding sites on the Illumina Infinium MethylationEPIC BeadChip. Genomics Data 9:22–24. https://doi.org/10.1016/j.gdata.2016.05.012

105. Zhou W, Laird PW, Shen H (2017) Comprehensive characterization, annotation and innovative use of Infinium DNA methylation BeadChip probes. Nucleic Acids Res 45:e22. https://doi.org/10.1093/nar/gkw967

106. Nordlund J, Bäcklin CL, Wahlberg P, Busche S, Berglund EC, Eloranta M-L, Flaegstad T, Forestier E, Frost B-M, Harila-Saari A, Heyman M, Jónsson ÓG, Larsson R, Palle J, Rönnblom L, Schmiegelow K, Sinnett D, Söderhäll S, Pastinen T, Gustafsson MG, Lönnerholm G, Syvänen A-C (2013) Genome-wide signatures of differential DNA methylation in pediatric acute lymphoblastic leukemia. Genome Biol 14:r105. https://doi.org/10.1186/gb-2013-14-9-r105

107. Stewart GB, Altman DG, Askie LM, Duley L, Simmonds MC, Stewart LA (2012) Statistical analysis of individual participant data meta-analyses: a comparison of methods and recommendations for practice. PLoS One 7:e46042. https://doi.org/10.1371/journal.pone.0046042

108. Smith-Warner SA, Spiegelman D, Ritz J, Albanes D, Beeson WL, Bernstein L, Berrino F, van den Brandt PA, Buring JE, Cho E, Colditz GA, Folsom AR, Freudenheim JL, Giovannucci E, Goldbohm RA, Graham S, Harnack L, Horn-Ross PL, Krogh V, Leitzmann MF, McCullough ML, Miller AB, Rodriguez C, Rohan TE, Schatzkin A, Shore R, Virtanen M, Willett WC, Wolk A, Zeleniuch-Jacquotte A, Zhang SM, Hunter DJ (2006) Methods for pooling results of epidemiologic studies: the Pooling Project of Prospective Studies of Diet and Cancer. Am J Epidemiol 163:1053–1064. https://doi.org/10.1093/aje/kwj127

109. Niu L, Xu Z, Taylor JA (2016) RCP: a novel probe design bias correction method for Illumina Methylation BeadChip. Bioinformatics 32:2659–2663. https://doi.org/10.1093/bioinformatics/btw285

110. Touleimat N, Tost J (2012) Complete pipeline for Infinium(®) Human Methylation 450K BeadChip data processing using subset quantile normalization for accurate DNA methylation estimation. Epigenomics 4:325–341. https://doi.org/10.2217/epi.12.21

111. Benjamini Y, Hochberg Y (1995) Controlling the False Discovery Rate: A Practical and Powerful Approach to Multiple Testing. Journal of the Royal Statistical Society Series B (Methodological) 57:289–300

112. Borisov V, Leemann T, Seßler K, Haug J, Pawelczyk M, Kasneci G (2022) Deep Neural Networks and Tabular Data: A Survey. arXiv:211001889 [cs]

113. Friedman J (2001) Greedy Function Approximation: A Gradient Boosting Machine. The Annals of Statistics 20:1189–1232. https://doi.org/10.1214/aos/1013203451

114. Zhao Y, Chetty G, Tran D (2019) Deep Learning with XGBoost for Real Estate Appraisal. In: 2019 IEEE Symposium Series on Computational Intelligence (SSCI). pp 1396–1401

115. Santhanam R, Uzir N, Raman S, Banerjee S (2017) Experimenting XGBoost Algorithm for Prediction and Classification of Different Datasets

116. Kingma DP, Ba J (2017) Adam: A Method for Stochastic Optimization. arXiv:14126980 [cs]

117. Little RJA, Rubin DB (2020) Statistical analysis with missing data, Third edition. Wiley, Hoboken, NJ

118. Bennett DA (2001) How can I deal with missing data in my study? Aust N Z J Public Health 25:464–469

119. Jerez JM, Molina I, García-Laencina PJ, Alba E, Ribelles N, Martín M, Franco L (2010) Missing data imputation using statistical and machine learning methods in a real breast cancer problem. Artificial Intelligence in Medicine 50:105–115. https://doi.org/10.1016/j.artmed.2010.05.002

120. Khan SI, Hoque ASML (2020) SICE: an improved missing data imputation technique. J Big Data 7:37. https://doi.org/10.1186/s40537-020-00313-w

121. Lin W-C, Tsai C-F (2020) Missing value imputation: a review and analysis of the literature (2006–2017). Artif Intell Rev 53:1487–1509. https://doi.org/10.1007/s10462-019-09709-4

122. Rubin LH, Witkiewitz K, Andre JS, Reilly S (2007) Methods for Handling Missing Data in the Behavioral Neurosciences: Don’t Throw the Baby Rat out with the Bath Water. J Undergrad Neurosci Educ 5:A71–77

123. Delalleau O, Courville A, Bengio Y (2018) Efficient EM Training of Gaussian Mixtures with Missing Data. arXiv:12090521 [cs, stat]

124. Andridge RR, Little RJA (2010) A Review of Hot Deck Imputation for Survey Non-response. International Statistical Review 78:40–64. https://doi.org/10.1111/j.1751-5823.2010.00103.x

125. Cheema JR (2014) A Review of Missing Data Handling Methods in Education Research. Review of Educational Research 84:487–508. https://doi.org/10.3102/0034654314532697

126. Jonsson P, Wohlin C (2004) An evaluation of k-nearest neighbour imputation using likert data. In: 10th International Symposium on Software Metrics, 2004. Proceedings. IEEE, Chicago, IL, USA, pp 108–118

127. Maillo J, Ramírez S, Triguero I, Herrera F (2017) kNN-IS: An Iterative Spark-based design of the k-Nearest Neighbors classifier for big data. Knowledge-Based Systems 117:3–15. https://doi.org/10.1016/j.knosys.2016.06.012

128. Amirteimoori A, Kordrostami S (2010) A Euclidean distance-based measure of efficiency in data envelopment analysis. Optimization 59:985–996. https://doi.org/10.1080/02331930902878333

129. Beretta L, Santaniello A (2016) Nearest neighbor imputation algorithms: a critical evaluation. BMC Medical Informatics and Decision Making 16:74. https://doi.org/10.1186/s12911-016-0318-z

130. Acuña E, Rodriguez C (2004) The Treatment of Missing Values and its Effect on Classifier Accuracy. In: Banks D, McMorris FR, Arabie P, Gaul W (eds) Classification, Clustering, and Data Mining Applications. Springer, Berlin, Heidelberg, pp 639–647

131. Lee JY, Styczynski MP (2018) NS-kNN: a modified k-nearest neighbors approach for imputing metabolomics data. Metabolomics 14:153. https://doi.org/10.1007/s11306-018-1451-8

132. Sun B, Ma L, Cheng W, Wen W, Goswami P, Bai G (2017) An improved k-nearest neighbours method for traffic time series imputation. In: 2017 Chinese Automation Congress (CAC). pp 7346–7351

133. Cheng D, Zhang S, Deng Z, Zhu Y, Zong M (2014) kNN Algorithm with Data-Driven k Value. In: Luo X, Yu JX, Li Z (eds) Advanced Data Mining and Applications. Springer International Publishing, Cham, pp 499–512

134. Murti DMP, Pujianto U, Wibawa AP, Akbar MI (2019) K-Nearest Neighbor (K-NN) based Missing Data Imputation. In: 2019 5th International Conference on Science in Information Technology (ICSITech). pp 83–88

135. Huang J, Keung JW, Sarro F, Li Y-F, Yu YT, Chan WK, Sun H (2017) Cross-validation based K nearest neighbor imputation for software quality datasets: An empirical study. Journal of Systems and Software 132:226–252. https://doi.org/10.1016/j.jss.2017.07.012

136. Zhu M, Cheng X (2015) Iterative KNN imputation based on GRA for missing values in TPLMS. In: 2015 4th International Conference on Computer Science and Network Technology (ICCSNT). pp 94–99

137. Zhang S, Li X, Zong M, Zhu X, Cheng D (2017) Learning *k* for kNN Classification. ACM Trans Intell Syst Technol 8:43:1-43:19. https://doi.org/10.1145/2990508

138. Samek W, Wiegand T, Müller K-R (2017) Explainable Artificial Intelligence: Understanding, Visualizing and Interpreting Deep Learning Models. arXiv:170808296 [cs, stat]

139. Caruana R, Lou Y, Gehrke J, Koch P, Sturm M, Elhadad N (2015) Intelligible Models for HealthCare: Predicting Pneumonia Risk and Hospital 30-day Readmission. In: Proceedings of the 21th ACM SIGKDD International Conference on Knowledge Discovery and Data Mining. Association for Computing Machinery, New York, NY, USA, pp 1721–1730

140. Lapuschkin S, Binder A, Montavon G, Muller KR, Samek W (2016) Analyzing Classifiers: 29th IEEE Conference on Computer Vision and Pattern Recognition, CVPR 2016. Proceedings - 29th IEEE Conference on Computer Vision and Pattern Recognition, CVPR 2016 2912–2920. https://doi.org/10.1109/CVPR.2016.318

141. Arras L, Horn F, Montavon G, Müller K-R, Samek W (2016) Explaining Predictions of Non-Linear Classifiers in NLP. In: Proceedings of the 1st Workshop on Representation Learning for NLP. Association for Computational Linguistics, Berlin, Germany, pp 1–7

142. Arras L, Horn F, Montavon G, Müller K-R, Samek W (2017) “What is relevant in a text document?”: An interpretable machine learning approach. PLOS ONE 12:e0181142. https://doi.org/10.1371/journal.pone.0181142

143. Schütt KT, Arbabzadah F, Chmiela S, Müller KR, Tkatchenko A (2017) Quantum-chemical insights from deep tensor neural networks. Nat Commun 8:13890. https://doi.org/10.1038/ncomms13890

144. Sturm I, Lapuschkin S, Samek W, Müller K-R (2016) Interpretable deep neural networks for single-trial EEG classification. Journal of Neuroscience Methods 274:141–145. https://doi.org/10.1016/j.jneumeth.2016.10.008

145. Lipovetsky S, Conklin M (2001) Analysis of regression in game theory approach. Applied Stochastic Models in Business and Industry 17:319–330. https://doi.org/10.1002/asmb.446

